# Lifelong Invertebrate Chordate Central Nervous System Regeneration is Driven by a Migrating Pool of Conserved Neural Stem Cells program

**DOI:** 10.64898/2026.09.11.750342

**Authors:** Chiara Anselmi, Tom Levy, Jiamu “Chester” Yu, Katherine J. Ishizuka, Karla J. Palmeri, Daniel Dan Liu, Rahul Sinha, Yotam Voskoboynik, Clelia Corridori, Norma F. Neff, Angela M. Detweiler, Irving L. Weissman, Graziano Martello, Ayelet Voskoboynik

**Author notes:** Equal senior contribution (Graziano Martello, Ayelet Voskoboynik).

## Abstract

Adult central nervous systems (CNS) are generally constrained by limited cellular replacement. How regenerating organisms integrate structural maintenance with continuous, cyclic renewal remains unresolved. Here, we leverage the colonial chordate *Botryllus schlosseri* which completely resorbs and regenerates its entire CNS *de novo* weekly. FACS, Combining indexed single-cell sorting, Smart-seq3 sequencing, spatial mapping, and temporal deconvolution, our studies suggest that this continuous CNS renewal is mediated by an intrinsic pool of lineage-restricted neural stem cells. Single-cell analysis resolved a conserved *SOX2*/*NOTCH1*/*NEUROG1* prospective progenitor continuum alongside mature excitability-defined neurons. Brain aging drives a phase-specific desynchronization of this regenerative program, inducing early hyper-activation followed by a sudden collapse. Finally, syngeneic transplantation and *in vivo* tracking show that neural-complex-derived cells persist after vascular delivery and localize to developing bud and neural-associated territories, with injury-associated accumulation near the neural gland. This work establishes a spatiotemporal framework for lifelong chordate CNS renewal and aging.

**Highlights:**

1. Smart-seq3 atlas resolves 6 cell states in the regenerating chordate brain
2. Conserved *SOX2*^+^/*NOTCH1*^+^/*NEURO1*^+^ continuum drives adult neurogenesis.
3. Neural-complex small cells home to bud niches and integrate upon turnover
4. Brain aging stems from ancestral stem-cell exhaustion across cyclic renewal

**In Brief:** Anselmi et al. establish a spatiotemporal single-cell atlas of the cyclically regenerating *Botryllus schlosseri* central nervous system, identifying a conserved *SOX2*^+^/*NOTCH1*^+^/*NEURO1*^+^neural progenitor continuum. Transplanted neural-complex-derived cells persist in vivo and localize to developing bud and neural-associated territories, while lifelong brain aging is driven by the cumulative exhaustion of this ancestral stem cell pool.

## INTRODUCTION

Adult nervous systems are generally defined by strict constraints on cellular replacement, continually balancing circuit maintenance against the progressive threat of degeneration. In vertebrates, adult neurogenesis is spatially restricted, neuronal loss is often only partially compensated, and aging and neurodegenerative disease progressively compromise both neural cell populations and their supporting niches (Terreros-Roncal et al., 2021;). In mammals, endogenous adult neurogenesis is restricted to specialized niches, such as the subgranular zone of the dentate gyrus and the subventricular zone, where neural stem cells (NSCs) remain responsive to environmental cues, physical exercise, and cognitive enrichment (Codega et al., 2014; Eriksson et al., 1998; Gage, 2000; van Praag et al., 1999). However, high-resolution single-cell multi-omic profiling reveals a severe physiological bottleneck: the vast majority of mammalian NSCs reside in a state of deep, cell-cycle-arrested quiescence (Audesse and Webb, 2020; Codega et al., 2014). Prospective isolation and single-cell profiling across the human lifespan further demonstrate that postnatal neural stem and progenitor cell lineages undergo an exponential decline in baseline self-renewal and activation capacity following infancy (Liu et al., 2023; Liu et al., 2026). Spatial transcriptomics indicates that during aging, these specialized neurogenic niches become progressively locked within an inactive, glial-and ependymal-like microenvironment, trapping non-dividing stem cells in dormancy and sharply curtailing endogenous repair (Maimon et al., 2024; Terreros-Roncal et al., 2021). The extreme rarity of active NSCs, the difficulty of tracking stem cell lineages *in vivo*, and the structural complexity of the mammalian brain present fundamental challenges for deciphering the universal principles governing neural renewal, niche maintenance, and degeneration.

In this context, tunicates provide an important comparative perspective. As marine invertebrate chordates, tunicates represent the sister group to vertebrates and occupy a key phylogenetic position for understanding the evolutionary origins of chordate nervous systems (Delsuc et al., 2018; Lemaire, 2011; Nanglu et al., 2023). Tunicates exhibit developmental and regenerative plasticities that contrast sharply with vertebrate constraints, serving as powerful systems to investigate how conserved neural gene programs, typically restricted to embryonic stages in vertebrates, can be recurrently redeployed during adult tissue renewal.

Among chordates, the colonial ascidian *B. schlosseri* provides an unparalleled *in vivo* platform for dissecting the cellular and molecular dynamics of lifelong neural renewal. A single *B. schlosseri* colony consists of repeating adult units, called zooids alongside developing buds at precise maturation states interconnected by a shared vascular network (Figure S0). Each zooid contains its own centralized and peripheral nervous system. Colonies develop through lifelong, synchronized blastogenetic cycles in which all adult zooids are simultaneously resorbed and replaced by their maturing buds (Kowarsky et al., 2021). During each weekly takeover phase, the mature central nervous system degenerates completely, adult organs are resorbed while a replacement CNS differentiates *de novo* within the next-generation bud (Figure S0A). Because *B. schlosseri* colonies can persist for decades, accumulating defined physiological and behavioral decline over chronological time (more than 20 years) (Anselmi et al., 2022; Bocci et al., 2024; Domen et al., 2026), this system bridges the gap between cyclical programmed degeneration and organismal aging. Our previous studies have shown that both cyclical turnover and long-term aging are characterized by progressive neuronal loss, compromised siphon-contraction reflexes, and the up regulation of conserved mammalian cell-death and neurodegeneration pathways (Anselmi et al., 2022; Anselmi et al., 2023; Bocci et al., 2024).

The adult tunicate CNS, termed the neural complex, is a composite structure that includes the cerebral ganglion (brain) and the adjacent neural gland complex (Braun and Stach, 2019; Kowarsky et al., 2021; Lucia Manni and Roberta Pennati, 2015) (Figures 1C and S0C). In *B. schlosseri*, this complex develops via two distinct developmental processes: embryogenesis and asexual blastogenesis (Kowarsky et al., 2021). Embryogenesis yields a specialized larval nervous system required for swimming, sensory orientation, and settlement, alongside a juvenile cerebral ganglion, that persists through metamorphosis into the founder oozooid (Anselmi et al., 2025). During subsequent blastogenic generations, the adult-type CNS is regenerated *de novo.* Histological studies suggest that neurogenesis initiates from a transient, dorsal epithelial thickening, the dorsal tube, from which cells delaminate and differentiate to generate both the cerebral ganglion and the neural gland complex (Burighel et al., 1998; Prünster et al., 2019). Notably, this dorsal neurogenic structure partially re-engages the anteroposterior patterning programs of the embryonic neural tube, suggesting that embryonic developmental cassettes are repeatedly co-opted during adult blastogenesis (Prünster et al., 2019). This shared origin provides a strong anatomical and developmental rationale to evaluate the neural-complex as an integrated functional unit rather than examining the cerebral ganglion alone.

**Figure 1.**
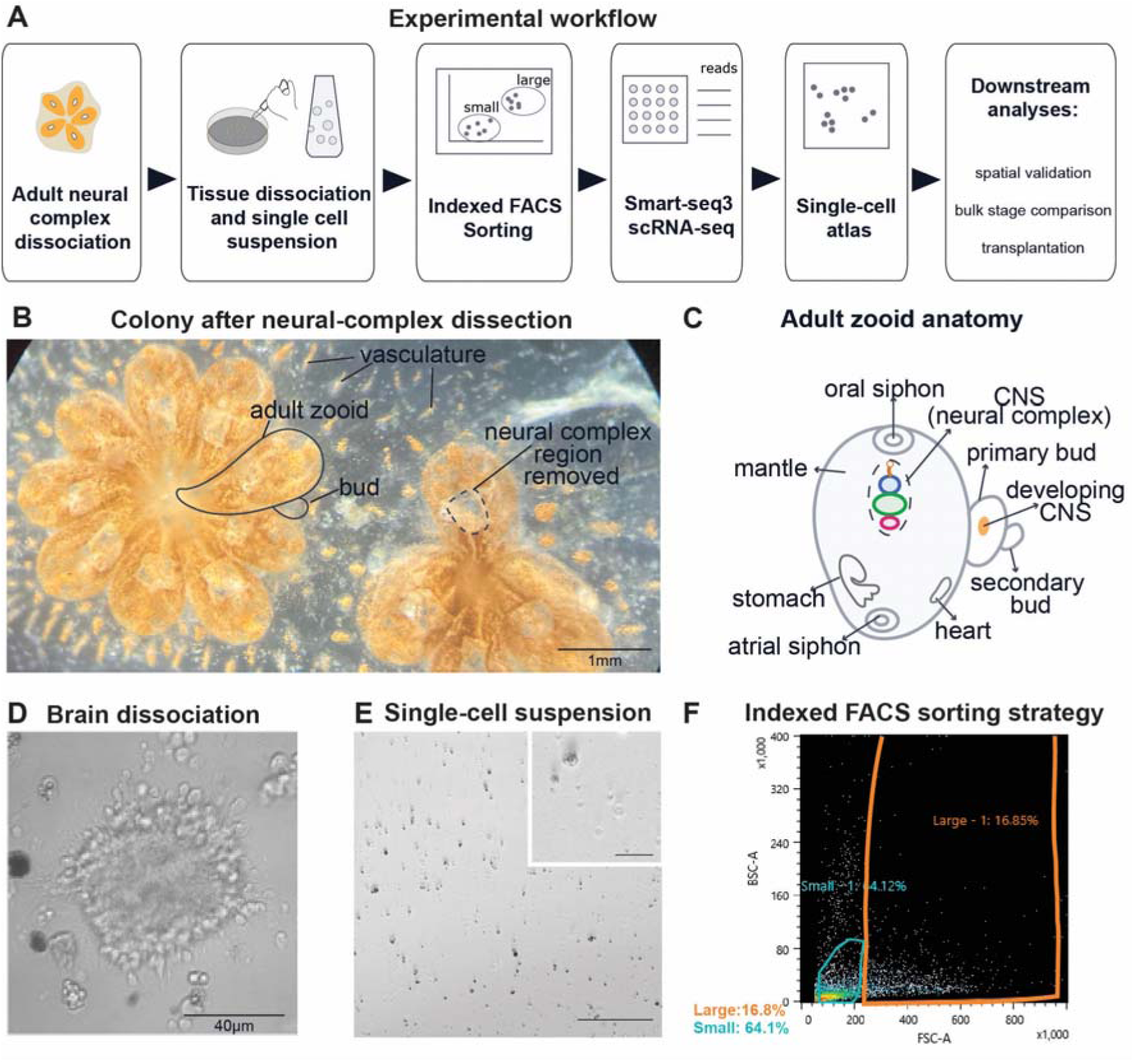
Experimental workflow and index-sorting pipeline for single-cell transcriptomic analysis of the adult *Botryllus schlosseri* brain. (A) Schematic overview of the experimental pipeline, including adult brain dissection, enzymatic and mechanical dissociation, single-cell suspension preparation, and index-based fluorescence-activated cell sorting (FACS) into 96-well plates for single-cell RNA sequencing. The resulting dataset was used to define neural-complex-associated cell states and to guide downstream spatial and comparative analyses. (B) Representative *B. schlosseri* colony attached to a substrate after removal of neural-complex regions from multiple adult zooids. *B. schlosseri* is a colonial tunicate in which clonally related zooids are embedded in a common tunic and connected by an extracorporeal blood vasculature. Each adult zooid contains its own central nervous system. The dashed outline highlights one representative adult zooid, while the dissected regions visible across the colony correspond to the areas containing the neural complexes that were removed from individual zooids for downstream single-cell profiling. The transparent body and shared vasculature of *B. schlosseri* make it suitable for live observation, dissection, and injection-based experimental manipulation. (C) Schematic overview of an adult zooid showing the position of the CNS between the oral and atrial siphons. The neural complex includes the cerebral ganglion/brain (green), neural gland (blue), ciliated duct and dorsal tubercle (orange), and finally the dorsal organ (pink). The brain sends nerves to the major organs and effector structures of the zooid body. The primary and secondary buds contain developing nervous systems. During the weekly blastogenetic cycle, degeneration of the adult brain occurs in parallel with continued neurogenesis in the bud generation that will replace the adult zooid. (D) Representative image of the dissected adult neural complex during dissociation. The neural complex appears as a compact central structure corresponding to the medulla, surrounded by a peripheral cortex. Axonal projections can be observed extending from the cortical region. (E) Dissociated neural-complex cells before FACS sorting. (F) FACS plot showing the gating strategy used to isolate dissociated cells for downstream single-cell analysis based on forward-and side-scatter properties, reflecting relative cell size and granularity.

Anatomically, the *B. schlosseri* cerebral ganglion exhibits a conserved chordate architecture: an outer cortex containing around 1000 neuronal somata, surrounding a dense, neurites central medulla. It is adjacent to the neural gland and the related dorsal organ (Anselmi et al., 2022; Mackie and Burighel, 2005). The neural gland is composed of large, vacuolated cells and is associated with the ciliated duct opening into the pharynx (Burighel et al., 1998). Despite its close anatomical association with the cerebral ganglion, the neural gland and related dorsal structures remain among the least understood components of the tunicate neural complex (Anselmi et al., 2022; Lucia Manni and Roberta Pennati, 2015). Whether these tissues participate directly in neural-complex remodeling, regeneration, or injury-associated responses has remained unresolved.

Multiple nerve roots emerge from the ganglion to interface with a widespread peripheral sensorimotor system activating the pharyngeal wall and branchial basket, which contains ciliated stigmata responsible for water flow (Mackie and Burighel, 2005). Importantly, primary sensory cells are distributed in peripheral epithelia, particularly around the oral and atrial siphons and within mantle-associated territories. These cells, characterized by a single cilium and direct axonal projections to the brain, function as mechanosensory neurons that relay environmental information to the cerebral ganglion (Anselmi et al., 2022; Anselmi et al., 2024). This centralized circuit regulates coordinated behavior, including spontaneous and stimulus-evoked siphon contractions, which fluctuate rhythmically across the weekly cycle in tandem with the maturation and regression of the adult zooid (Anselmi et al., 2022; Thompson et al., 2022; Zaniolo et al., 2002).

Beyond its capacity for complete neural regeneration, *B. schlosseri* offers a unique opportunity to ask whether neural renewal is coordinated with the broader stem-cell and niche biology of the colony. Previous work has shown that selected somatic and germline progenitor populations can circulate through the shared vasculature, localize to defined developmental niches, and contribute to tissue renewal in systems such as the endostyle, germline, cell islands, and peripheral ampullae (Laird et al., 2005; Levy et al., 2025; Rinkevich et al., 2013; Rosental et al., 2018; Vanni et al., 2023; Voskoboynik and Weissman, 2015; Voskoboynik et al., 2008).

However, despite extensive morphological descriptions, the cellular composition, transcriptional heterogeneity, and progenitor-associated programs accompanying lifelong nervous system replacement have remained undefined. Whether the *B. schlosseri* neural complex deploys conserved progenitor programs during cyclical renewal, how neural and non-neural cell states are spatially organized, and how these programs change across regenerative phases and aging remain fundamental open questions.

Here, we provide a comprehensive cellular, molecular, and spatial atlas of the regenerating adult tunicate nervous system across its complete life cycle. By integrating index-sorted Smart-seq3 single-cell RNA sequencing, multiplexed spatial mapping via whole-mount Hybridization Chain Reaction - Fluorescence In Situ Hybridization (HCR-FISH) and immunostaining, longitudinal bulk transcriptomics across blastogenetic and aging timepoints, and prospective *in vivo* cell transplantation assays, we resolve the spatiotemporal architecture of cyclical neural turnover. We find that the *B. schlosseri* neural complex contains distinct progenitor, neuronal, structural, and immune-associated cell states organized within specialized functional microenvironments. Progenitor populations actively express conserved chordate neurogenic programs and transition through distinct proliferative and differentiation states across the weekly cycle. Furthermore, *in vivo* tracking shows that neural-complex-derived cells can persist after transplantation and localize to developing bud and neural-complex-associated territories. Together, our results define conserved progenitor-like and neural programs associated with repeated CNS renewal and establish *B. schlosseri* as a tractable model for investigating adult neurogenesis, cyclical degeneration, and lifelong tissue maintenance.

## RESULTS

### A single-cell atlas resolves the cellular architecture of the adult *B. schlosseri* neural complex

To define the cellular composition of the adult *B. schlosseri* CNS (the neural complex), we generated a Smart-seq3 single-cell RNA-seq dataset from manually dissected neural-complex regions, comprising the cerebral ganglion and closely associated structures, the neural gland, ciliated duct, dorsal tubercle/dorsal organ region, and perineural margins (Figures 1A-C and S0C-D). Adult colonies were imaged before and after dissection to document the colonial context of the sampled tissue (Figure 1B).

Following dissociation, neural-complex-enriched cell suspensions were sorted by indexed FACS and processed for Smart-seq3 single-cell RNA sequencing (Figure 1A and 1D–F). After quality-control, 470 of 676 sorted cells, (69.5%) (Figure S1) and 17,118 genes were retained (Figure S1). Both *Uniform Manifold Approximation and Projection (*UMAP) and *t-distributed Stochastic Neighbor Embedding* (t-SNE) resolved six major transcriptionally distinct cell populations (Figures 2A and S2), which formed the basis for marker discovery, HCR-FISH validation, cell-cycle analysis, and bulk RNA-seq integration. Indexed FACS linked each cell’s physical properties to its transcriptomic identity: small and large-cell gates were biasedly distributed across UMAP space, indicating biologically distinct populations (Figures 1F and 2B). Cluster sizes ranged from abundant to rare, confirming capture of both major and minor cellular compartments (Figure 2C).

**Figure 2.**
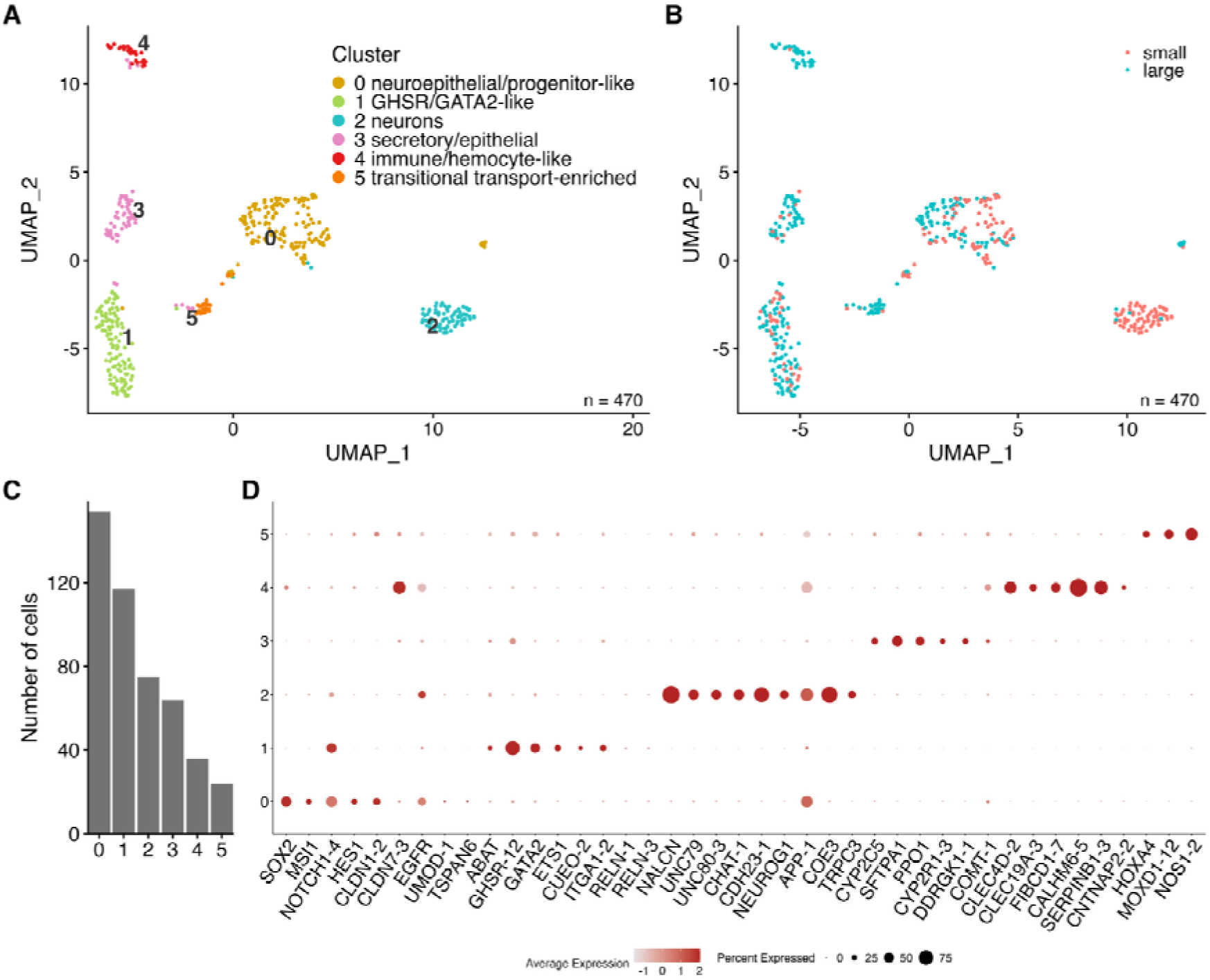
A single-cell transcriptomic atlas of the adult *Botryllus schlosseri* CNS reveal six distinct cellular clusters capturing neuronal, progenitor like, epithelial-like, immune like and transitional states. (A) UMAP representation of single cell transcriptomes resolving six transcriptionally distinct cellular states color-coded by unsupervised cluster assignment. (B) UMAP showing the distribution of sequenced cells according to their original FACS forward/side-scatter size gates (small versus large populations). (C) Bar plot showing the number of cells assigned to each cluster. (D) Dot plot of selected marker genes across clusters. Dot size indicates the percentage of cells expressing each gene, and color intensity indicates average scaled expression.

### Single-cell profiling defines progenitor, neuronal, metabolic-epithelial, and immune effector states

UMAP visualization resolved six transcriptionally distinct clusters within the adult *B. schlosseri* neural complex (Figure 2). Because the dissected region included the cerebral ganglion/brain together with closely associated neural-complex structures, including the neural gland and ciliated duct (Figure S0E), this transcriptional heterogeneity likely reflects both neural and brain-adjacent cellular compartments.

To assign cluster identities, we used updated gene models of the *B. schlosseri* genome (Levy et al., 2025), and an annotation workflow integrating differential marker analysis, curated anchor panels, and ontology enrichment (Figure S3 and Table S1-S2). Confidence was highest for clusters 0, 2, 4 and 5,whereas cluster 1 was interpreted cautiously as a regulatory state.

Cluster 0 displayed a distinct progenitor signature, enriched for core neural stem cell regulators (*SOX2*, *MSI1*, *NOTCH1*, *HES1*/*HES5*) and *CLDN* tight-junction genes, with gene ontology (GO) terms dominated by CNS development cell-cell adhesion and neurogenesis (Figure 2D).

Cluster 1 expressed the growth hormone secretagogue receptor *GHSR*, transcription factors *GATA2* and *ETS*, alongside metabolic regulators *ODC1*, and *ABAT* (GABA transaminase), suggesting possible neuroendocrine-like regulatory features.

Cluster 2 represented the differentiated neuronal core, with coordinated upregulation of genes involved in neuronal excitability (*NALCN*, *UNC79*, and *UNC80*), cholinergic transmission (*CHAT*, *ACHE*), and synaptic machinery (*NRXN*, *NLGN*, *NCAM*, *SNAP25*, *SYT1*), with GO terms dominated by synaptic organization, and ion/ neurotransmitter transport.

Cluster 3 is strongly enriched for metabolic/secretory genes (*CYP2C5*, *CYP2R1*, *SFTPA1*, *PPO1*, *DDRGK1*), matching the adjacent neuroendocrine neural gland.

Cluster 4 expressed innate recognition receptors and defense-related genes (*CLEC4D*, *CLEC19A*, *FIBCD1*, *CALHM6*, *SERPINB1*), supporting an immune/hemocyte-like effector identity participating in tissue remodeling.

Cluster 5 is co-enriched of homeobox patterning factors (*HOXA4*), solute transporters (*SLC5A6*, *SLC5A8*), *MOXD1*, and *NOS1*, consistent with a transitional, lineage-primed, or transport-active state within the broader neural-complex microenvironment.

Together, these annotations demonstrate that the adult *B. schlosseri*neural complex is not a uniform ganglion, but a structured territory integrating progenitor-like, neuronal, epithelial, immune-like, and transitional states, with clearest separation observed between the *SOX2*/*NOTCH*/*HES* progenitor program (cluster 0) and the *NALCN*/*CHAT*/*ACHE* neuronal program (cluster 2) (Figures 2D and S3).

### HCR-FISH maps single-cell states onto brain, neural-gland, mantle, and sensory territories

Whole-mount HCR-FISH of candidate markers identified from the scRNA-seq dataset, paired with UMAP feature plots, spatially anchored the atlas across colonies from Venice Lagoon and Monterey Bay (Figures 3 and S4, Tables S3 and S4). Negative-control samples processed in the absence of target probes showed minimal background fluorescence across adult and developing neural-complex-associated territories (Figure S5). Markers resolved into three primary spatial compartments: (i) Neuronal markers localized within the adult cerebral ganglion (brain), (ii) Progenitor-like and regulatory markers located around the brain and in the surrounding mantle tissue, and (iii) neural gland/ sensory epithelial.

**Figure 3.**
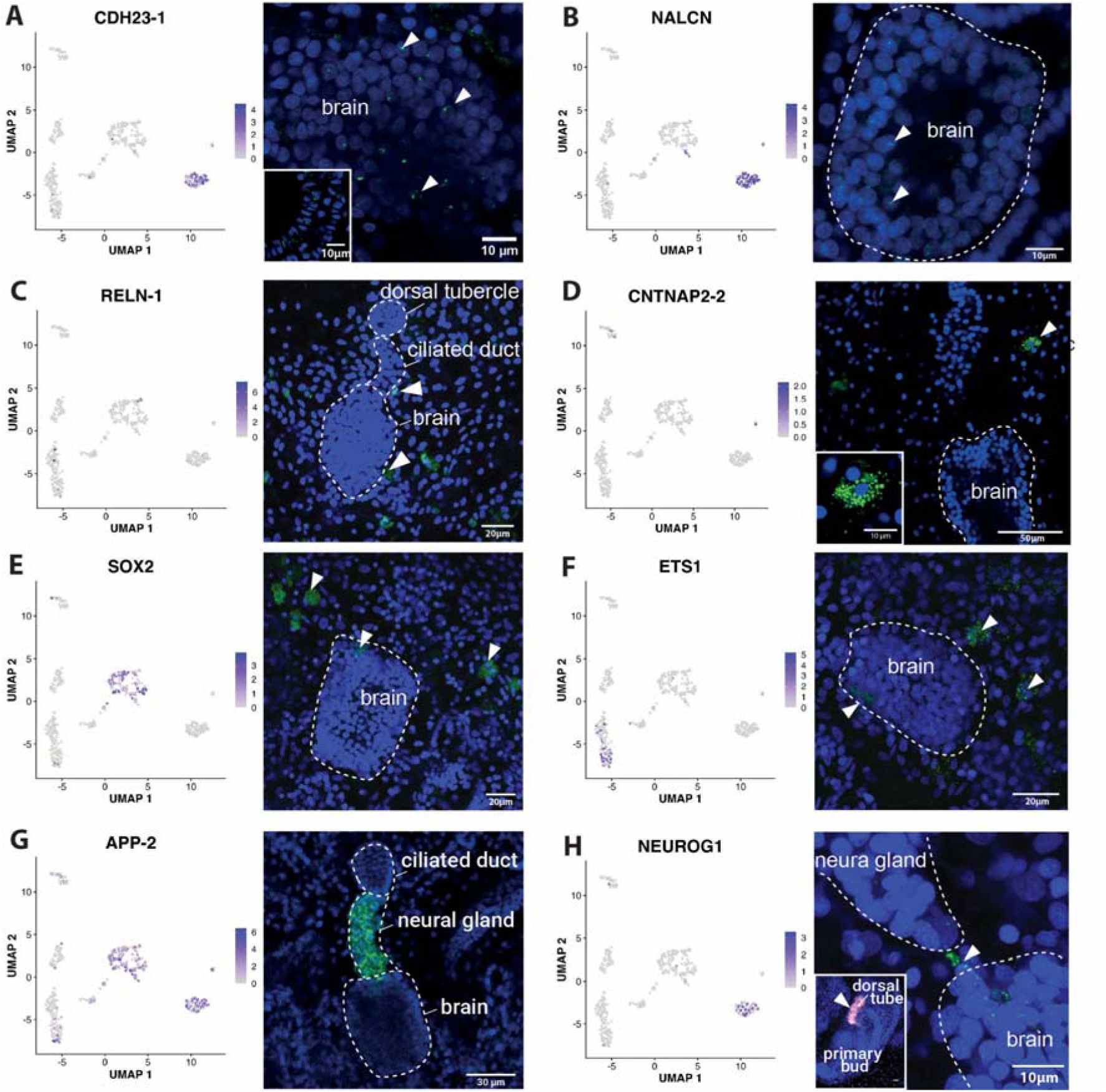
Spatial mapping of single-cell transcriptomic states via multiplexed HCR-FISH reveals the anatomical compartmentalization of the *Botryllus schlosseri* neural complex. Each panel combines a UMAP feature plot from the Smart-seq3 single-cell atlas, shown on the left, with HCR-FISH spatial validation in adult zooids or developing buds, shown on the right. Feature plots show normalized expression of selected marker genes across the adult *B. schlosseri* neural-complex atlas, while HCR-FISH identifies the anatomical localization of the corresponding transcripts in the neural complex and associated tissues. Nuclei are counterstained with DAPI in blue; HCR-FISH signal is shown in green, except for *NEUROG1* in the primary bud inset, where signal is shown in magenta. Dashed outlines indicate the brain/cerebral ganglion, neural gland, ciliated duct, dorsal tubercle, or developing bud neural territories when visible. Arrowheads indicate HCR-positive cells or expression domains. Scale bars are indicated in each panel. (A) CDH23*-1* is enriched in the neuronal cluster on the UMAP and is detected by HCR-FISH in cells associated with the adult brain region; the inset highlights signals in peripheral sensory cells in the oral siphon. (B) *NALCN* shows expression in the neuronal cluster and localizes to cells within the adult brain. (C) *RELN-1* is detected in sparse cells associated with the dorsal mantle and ciliated duct/neural-complex region. (D) *CNTNAP2-2* shows restricted expression in the single-cell atlas and HCR-FISH signal in discrete cells near the adult brain region, with higher-magnification inset showing clustered positive signal. (E) *SOX2* is primarily enriched in the neuroepithelial/progenitor-like cluster and labels cells around or adjacent to the adult brain. (F) *ETS1* marks scattered cells surrounding the brain and in adjacent body-wall territories. (G) *APP-2* is expressed across neural-complex-associated cells in the atlas and shows strong spatial signals in the neural gland region adjacent to the brain. (H) *NEUROG1* is enriched in the neuronal cluster and labels rare cells associated with the neural-complex region, including cells near the neural gland and brain; the inset highlights signals (in orange) in the dorsal tube of the primary bud.

We first examined markers associated with the neuronal compartment of the cerebral ganglion. A primary cohort of candidate markers confirmed the presence of a *bona fide* central neuronal compartment within the adult cerebral ganglion (brain). *CDH23* was robustly expressed in the adult brain cortex, where numerous neuronal somata displayed distinct punctate nuclear and perinuclear signaling (Figures 3A and S4A). In addition, *CDH23* was highly expressed in the inner epithelium of the oral siphon, a region enriched with primary mechanosensory cells, while remaining absent in developing buds or other mantle territories (Figure 3A). *NALCN*, associated with neuronal excitability, showed a spatial pattern consistent with the neuronal cluster identified by scRNA-seq. *NALCN* transcripts were detected within the primary bud brain with intense signals observed in scattered adult mantle cells near the neural complex (Figures 3B and S4A). Its accessory subunits *UNC80* and *UNC79* showed a more brain-centered expression pattern: *UNC80* was detected across both adult and primary buds brain, whereas *UNC79* showed a more centrally enriched pattern, largely restricted to the adult brain (Figure S4A). *TRPC3* was also detected within the adult brain, with signal distributed between nuclei and consistent with cytoplasmic expression in brain-associated cells; additional scattered mantle-associated cells were also labeled (Figure S4A). Additional neuronal support came from *COE3*, which was detected within the brain and in brain-associated cells (Figure S4A). Together, these markers spatially validate the neuronal and excitability programs of cluster 2.

A second group of markers identified brain-adjacent progenitor-like and regulatory populations. *RELN* showed a highly restricted and reproducible pattern across three independent colonies. In all adult zooids examined, *RELN* was detected in isolated distinct cells within the dorsal mantle and in scattered cells near the oral siphon. In addition, a few neurons at the periphery of the brain cortex were positive (Figures 3C and S4B). No *RELN* expression was detected in buds or other adult tissues. CNTNAP2, a neurexin-family adhesion molecule, was consistently detected in cells close to the adult brain, showing both nuclear-associated labeling and strong concentrations of small marked areas in the cytoplasm (Figures 3D and S4C). *CNTNAP2* signal was also observed in the developing primary bud brain, whereas no other adult tissue was labeled (Figure S4C). TSPAN6, a tetraspanin, showed consistent signal in the outer cortex of the brain and in isolated mantle cells, with no detectable expression in buds (Figure S4D). The pluripotent/multipotent marker SOX2 was detected in specific cells adjacent to the adult brain and in cells within the brain itself, with no detectable expression in buds or in other adult tissues (Figures 3E and S4D). This pattern is consistent with a brain-adjacent, non-uniformly distributed population rather than broad expression throughout the ganglion. Finally, the transcription factor ETS1 marked scattered cells surrounding the brain and in the body wall (Figures 3F and S4B). Together, these data define a broader, specialized brain-adjacent compartment composed of progenitor-like, regulatory, and mantle-associated cells strategically positioned around the adult neural complex.

A third group of markers mapped neural-gland-associated, developing, and peripheral sensory or epithelial territories. In adult zooids, APP was consistently detected throughout the neural complex, showing a prominent enrichment in the neural gland and ciliated duct region, alongside weaker expression in the brain cortex (Figures 3G and S4E). This gland-associated enrichment was particularly evident in stage A adults, where APP labeling was stronger in the neural gland/ciliated duct than in the cerebral ganglion. APP expression was also observed in primary sensory cells distributed in the mantle and around the oral siphon. Across later stages, *APP* remained detectable in adult neural complexes undergoing regression during takeover, while signal was also present in the developing neural complex of primary buds that will replace the adult zooids. In stage C2 colonies, *APP* signal was detected in the neural complex of adult zooids as well as in primary and secondary buds. These dynamic expression profiles indicate that *APP* expression is not confined to differentiated adult brain neurons, but instead marks neural-gland-associated, sensory, degenerating, and developing neural-complex territories across the blastogenetic cycle (Figures 3G and S4E). In contrast, *ITIH4* expression was localized exclusively to the neural gland, with no detectable signal in the brain, indicating that this marker identifies a gland-associated rather than neuronal territory (Figure S4D). Within the scRNA-seq dataset *ITIH4* was detected in a small subset of cells and restricted to cluster 0. Thus, *ITIH4* was retained primarily as an anatomically informative HCR marker of the neural gland region, rather than as a major transcript used for cluster annotation. *UMOD* showed a strikingly peripheral pattern. In all adult zooids examined, *UMOD* signal was excluded from the central neural tissues and buds, mapping exclusively to the inner and outer epidermis of the oral siphon and the terminal cloacal languet converging centrally to form the atrial siphon (Figure S4D). Thus, *UMOD* marks peripheral epithelial or sensory-associated territories rather than central neural tissue. The conserved proneural factor *NEUROG1*, which drives neural commitment in vertebrates, (Ma et al., 1996) showed a restricted and stage-dependent pattern associated with neural-complex development (Figures 3H and S4F). *NEUROG1* prominently marked the dorsal tube of the primary bud, consistent with the transient neurogenic territory that gives rise to the neural gland and cerebral ganglion. Before takeover, the *NEUROG1* signal In primary buds was detected within the developing neural complex, with stronger labeling in the neural gland region than in the developing brain. This localized tracking highlights the role of *NEUROG1* in specifying early neural identity and coordinating the downstream neurogenic cascade required to build the organ *de novo* during budding. In fully mature adult zooids, where neurogenesis is largely complete, *NEUROG1* expression dropped significantly, limited to rare cells within or adjacent to the cerebral ganglion, including cells located between the brain and neural gland.

Collectively, these spatial validation data demonstrate that the single cell atlas successfully decodes the dissected anatomy, spanning central cerebral ganglion neurons, neural gland, developing bud neurogenic zones, and peripheral sensory-epithelia (Figure S4G).

### Cell-cycle scoring, trajectory inference, and PH3 staining reveal dynamic neurogenic states

Because the single-cell atlas identified progenitor-like, proneural, and transitional transcriptional states within the adult neural complex, we next investigated whether these populations were associated with active proliferation. We computationally inferred the cell-cycle phase from the single-cell transcriptomes using expression-based cell-cycle scoring and complemented this with phospho-histone H3 (PH3) immunostaining across the blastogenetic cycle (Figures S6A, S6B and S7, Table S3).

Cell-cycle scoring was not uniformly distributed across clusters (Figure S6A). Clusters 1 and 2 had the highest S/G2/M fractions and cluster 4 (immune effectors) showed the highest G1 fraction (Figure S6A). Examining cell-cycle state relative to *SOX2* and *NEUROG1* revealed distinct kinetic behaviours. *SOX2*-positive cells (predominantly in progenitor-like cluster 0), displayed a mixed G1 and S/G2/M profile, consistent with a heterogeneous progenitor pool containing both cycling and non-cycling cells. In contrast, *NEUROG1*-positive cells, (concentrated in neuronal cluster 2) were predominantly assigned to S/G2/M phases, suggesting that *NEUROG1* expression in the adult *B. schlosseri* CNS marks an actively proliferative or recently cycling proneural/neuronal state rather than exclusively a terminally differentiated neuronal population. *SOX2*/*NEUROG1* double-positive cells were extremely rare. Slingshot trajectory inference independently connected clusters 0 and 2 within the unrooted transcriptional topology. When cluster 0 was specified as a biologically informed starting state, the inferred 0–2 axis was characterized by *SOX2*/*MSI1*/*HES*/*NOTCH*-associated expression toward earlier states and *NEUROG1*/*NALCN*/*UNC79*/*UNC80*/*CHAT*/*ACHE* expression toward later states (Figure S6B). Within this axis, *SOX2*-positive cells decreased in the terminal pseudotime quartile of cluster 0, whereas *NEUROG1*-positive cells were depleted in the earliest quartile of cluster 2 and enriched thereafter, supporting an ordered transcriptional relationship between progenitor-like and proneural/neuronal states without establishing direct lineage progression.

To spatially assess mitotic activity across the blastogenetic cycle, we performed whole-mount immunostaining for PH3 alongside alpha tubulin and DAPI across defined developmental stages using (Figure S7). In primary and secondary buds, PH3-positive cells were observed abundantly throughout all blastogenic stages, consistent with ongoing de novo organogenesis. In contrast, the adult neural complex showed a temporally and spatially restricted mitotic pattern. PH3-positive cells in adult zooids were detected mainly at early stages (Days 1-3) and were localized specifically to the dorsal organ and ciliated duct/neural gland regions, becoming virtually undetectable prior to takeover (Figure S7). Together, these analyses indicate that neurogenic proliferation is sustained continuously in developing buds and is restricted to early-cycle remodeling niches in adult zooids neural-complex-associated regions.

### Bulk brain RNA-seq reveals cyclic remodeling of single-cell-defined programs

To determine whether single-cell-defined transcriptional programs undergo coordinated shifts at the whole-tissue level across the weekly cycle and and chronological aging, we generated a longitudinal bulk RNA-seq dataset from dissected *B. schlosseri* neural complexes across blastogenetic stages (Days 1-7) and colony lifespans (young: 47-243 days; old:3721-6845 days, Figures 4 and S8). Principal component analysis of young adult bulk transcriptomes revealed structured stage-associated separation, with late stages, (Days 5 and 7), diverging from early stages (Days 1 and 3) (Figure S8A).

**Figure 4.**
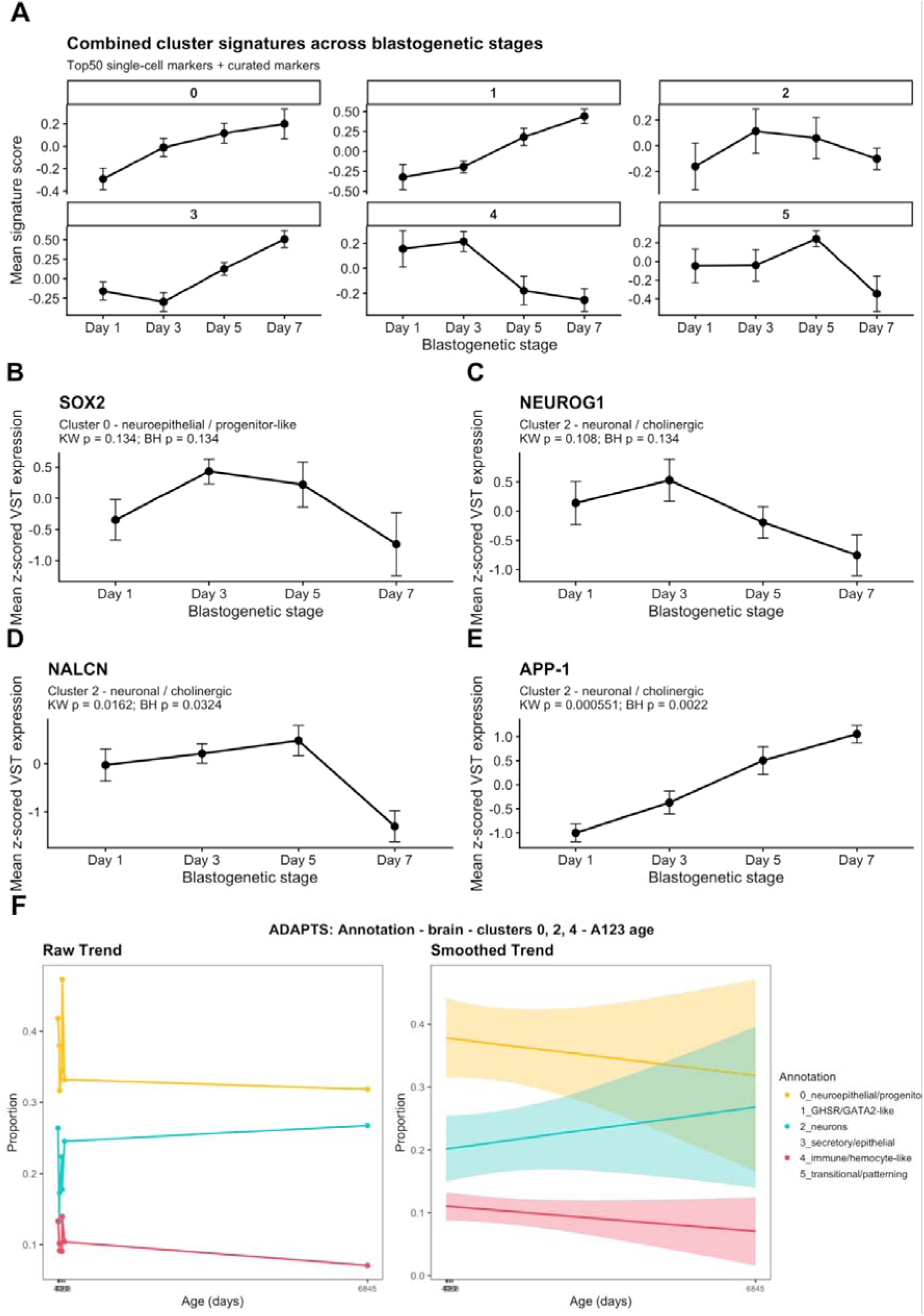
Bulk RNA-seq analysis identifies blastogenetic-stage-associated transcriptional dynamics in *Botryllus schlosseri* brains. (A) Combined cluster-associated signature scores across the blastogenetic cycle in young adult brains. Signature scores were calculated from bulk RNA-seq data using the mean z-scored VST expression of single-cell-derived Top 50 cluster markers together with manually curated marker genes. Samples are grouped as Day 1, Day 3, Day 5, and Day 7, corresponding to blastogenetic stages A, B, C, and D. Each panel represents one single-cell-derived cluster. Points show the mean signature score per stage and error bars indicate standard error of the mean. Cluster 0, corresponding to the neuroepithelial/progenitor-like program, increases toward later stages. Cluster 3 also increases strongly toward Day 7, whereas clusters 4 and 5 decrease at Day 7. (B–E) Stage-dependent expression of selected marker genes from biologically relevant cluster programs. Expression is shown as mean z-scored VST expression ± standard error across Day 1, Day 3, Day 5, and Day 7. (B) *SOX2*, associated with the neuroepithelial/progenitor-like cluster 0, peaks around Day 3-Day 5 and decreases by Day 7. (C) *NEUROG1*, associated with the neuronal/cholinergic cluster 2, follows a similar early-to-mid-stage pattern and decreases toward Day 7. (D) *NALCN*, also associated with cluster 2, increases toward Day 5 and sharply decreases at Day 7. (E) *APP-1*, associated with cluster 2, progressively increases from Day 1 to Day 7. Each panel reports the raw *p* value from the Kruskal–Wallis test and the corresponding *p* value after Benjamini–Hochberg adjustment. (F) ADAPTS-based deconvolution of bulk brain RNA-seq samples across age using the single-cell brain atlas as reference. Estimated proportions of clusters 0, 2, and 4 are shown across *B. schlosseri* samples collected at A1-A3 stages. The left panel shows raw estimated cluster proportions across age, while the right panel shows smoothed age-associated trends with confidence intervals. Colors indicate single-cell-derived cluster annotations.

Projecting scRNA-derived signature scores (mean z-scored VST expression of single-cell-derived marker genes, combining top cluster markers with curated marker genes) across the blastogenic cycle demonstrated that single-cell modules follow distinct temporal trajectories (Figure 4A). The neuroepithelial/progenitor-like signature (cluster 0) and the secretory/metabolic epithelial-like signature (cluster 3) increased progressively toward day 7 (Figure 4A). In contrast, the immune/hemocyte-like signature (cluster 4) and transitional/patterning-enriched signature (cluster 5) decreased towards Day 7. Progenitor associated *SOX2*, and proneural *NEUROG1* peaked during mid-cycle (Days 3-5) and declined toward day 7 (Figures 4B-4C). Neuronal excitability marker *NALCN* peaked at Day 5 before sharply dropping at Day 7 (Figure 4D). In contrast, *APP-1* showed a monotonic, progressive increase from Day 1 to Day 7, suggesting that selected neural-complex-associated genes remain elevated or increase during late-cycle remodeling (Figure 4E).

Intersecting DEseq2 likelihood ratio test (LRT) stage-regulated bulk genes with single-cell marker frameworks confirmed that whole tissue cyclical turnover reflects coordinated remodeling across neuronal, progenitor, epithelial and immune modules (Figure S8B). Because bulk RNA-seq measures the averaged transcriptional output of the entire neural complex, we next asked whether age-associated differences could reflect changes in the relative representation of the cell states identified by scRNA-seq. We therefore applied ADAPTS-based reference deconvolution, using the single-cell cluster expression profiles as a reference to estimate the relative contribution of each transcriptional program to the bulk transcriptomes. In Stage A (day 1) neural complexes, comparison of young and old colonies suggested age-related alterations in the estimated representation of progenitor-like (cluster 0), neuronal (cluster 2), and immune-associated (cluster 4) programs (Figure 4F). Together, these results are consistent with the possibility that chronological aging is associated with altered relative representation of neural-complex-associated transcriptional programs during early adult blastogenesis.

### Neural-complex-derived small cells localize to developing and neural-associated territories after transplantation

Previous studies in *B. schlosseri* have shown that selected somatic and germline progenitor populations can circulate through the shared vasculature, localize to defined developmental niches, and contribute to tissue renewal (Laird et al., 2005; Levy et al., 2025; Rinkevich et al., 2013; Rosental et al., 2018; Voskoboynik et al., 2008). This circulatory trafficking and targeted mobilization mirror hallmark properties of mammalian stem cells established by prospective isolation and *in vivo* tracking: hematopoietic stem cells constitutively enter the blood circulation to migrate and re-engraft unconditioned bone marrow niches (Wright et al., 2001), while prospectively isolated human neural stem cells (Liu et al., 2023; Uchida et al., 2000) actively migrate across parenchyma toward specialized neurogenic niches or undergo targeted long-distance migration to sites of CNS injury and ischemia (Kelly et al., 2004). We therefore asked whether cells isolated from the adult neural complex could persist after vascular transplantation and display distinct localization patterns within recipient colonies.

To test this, FACS-isolated small-and large-cell fractions from stage A (day 1) donor neural complexes were labelled with the lipophilic dye Vybrant DiD and microinjected into the extracorporeal vasculature of stage-matched isogeneic histocompatible recipient colonies, typically through proximal ampullae or blood vessels following an established cell-labelling and transplantation approach for *B. schlosseri* (Levy et al., 2025) (Figure 5A). Live *in vivo* imaging revealed that DiD-labelled small cells could be detected in adult-associated and bud-associated territories of recipient colonies.

**Figure 5.**
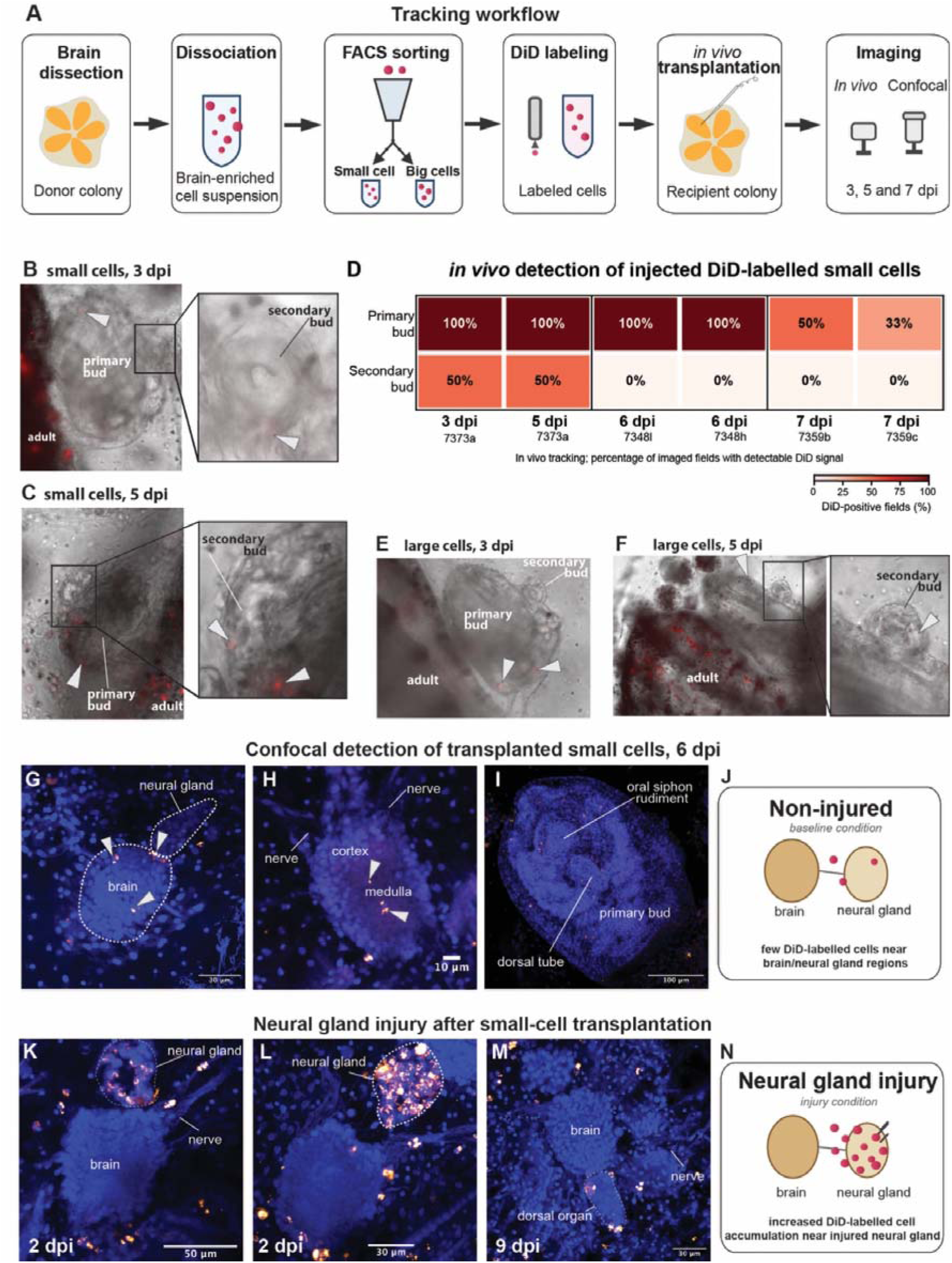
Neural-complex-derived cells are detected in adult and bud territories after transplantation and accumulate near the neural gland following injury. (A) Tracking workflow. Adult brains/neural complexes were dissected from donor colonies, dissociated into brain-enriched cell suspensions, separated by FACS into small-and large-cell fractions, labelled with DiD, transplanted into recipient colonies through the vasculature, and imaged *in vivo* or by confocal microscopy at the indicated days post injection (dpi). (B–C) Representative *in vivo* images of recipient colonies injected with DiD-labelled small cells at 3 dpi and 5 dpi. Labelled cells are detected in adult zooid regions and developing buds. Boxed regions are shown at higher magnification. (D) Heatmap summary of *in vivo* detection of injected DiD-labelled small cells in developing bud territories across individual recipient colonies and time points. Values indicate the percentage of imaged fields with detectable DiD signal in the primary bud and secondary bud compartments. Because the small-and large-cell fractions were not transplanted at equivalent cell numbers, quantitative summary is shown for the small-cell fraction only. Each column represents an individual recipient colony/subclone; the colony ID is shown below each column and the corresponding imaging time is indicated above in days post injection (dpi). (E–F) Representative *in vivo* images of recipient colonies injected with DiD-labelled large cells at 3 dpi and 5 dpi. Large-cell-derived DiD signal was detected less frequently than small-cell-derived signal. (G–I) Confocal detection of DiD-labelled small cells at 6 dpi in non-injured recipient colonies. Labelled cells are observed near the adult brain/neural gland region, within or near the brain cortex and medulla, along nerves, and in developing bud territories including the dorsal tube and oral siphon rudiment region. Lower-magnification views documenting signal specificity within the recipient zooid are provided in Figure S10. (J) Summary schematic of the non-injured condition, in which few DiD-labelled cells are detected near brain/neural gland-associated regions. (K–M) Confocal images following neural gland injury after transplantation of DiD-labelled small cells. Labelled cells accumulate near the neural gland, brain-associated region, nerves, and dorsal organ-associated territories at the indicated time points. Lower-magnification views documenting signal specificity within the recipient zooid are provided in Figure S10. (N) Summary schematic of the neural gland injury condition, showing increased DiD-labelled cell accumulation near the injured neural gland. DiD-labelled cells are shown in red/magenta and nuclei are counterstained with DAPI in blue. White arrowheads indicate DiD-labelled transplanted cells. Dashed outlines indicate anatomical regions. Scale bars are indicated in each panel.

At 3 days post injection (dpi), small-cell-derived DiD signal was detected in multiple recipient territories, including cell islands and developing buds (Figure 5B). In the recipients summarized in Figure 5D, primary-bud detection ranged from 33% to 100% of imaged fields, whereas secondary-bud detection ranged from 0% to 50% and was observed in a subset of recipients. Representative *in vivo* images of colonies transplanted with DiD-labelled large cells are shown in Figures 5E–5F, and additional localization patterns are provided in Figure S9.

To examine the anatomical localization of transplanted cells at higher resolution, recipient colonies were fixed after transplantation and analyzed by confocal microscopy. In non-injured recipients examined at 6 dpi, confocal z-stack imaging showed that DiD-labelled cells from the small-cell fraction were located within the adult brain volume of the replacement-generation brain, including cortical and medullary regions (Figures 5G–5I and S10). Labelled cells were also detected at the brain-neural gland interface, along nerve-associated structures, and in developing bud territories, including the dorsal tube and oral siphon rudiment region. Extended confocal datasets further documented labelled small cells within and around the adult brain, in primary and secondary bud-associated territories, and along neural-complex-associated structures (Figure S10). Sham-transplanted control colonies injected with the Vybrant DiD dye solution in the absence of cells, alongside untreated subclonal controls, showed no detectable background fluorescence within the neural complex or budding zones, supporting the interpretation that the observed fluorescence was associated with transplanted donor-derived cells rather than nonspecific background from the dye.

The 6 dpi time point is biologically informative because of the weekly blastogenetic cycle of *B. schlosseri* (Figures 5G-5J). By this time, the zooid generation present at the time of injection has progressed, and tissues that were developing buds during transplantation can contribute to the replacement adult generation. Therefore, detection of DiD-labelled cells within the adult brain at 6 dpi indicates that a subset of transplanted neural-complex-derived small cells can localize within neural tissue of the replacement generation. The z-stack localization within the adult brain volume shows that transplanted neural-complex-derived cells are not merely circulating or externally associated with the tissue, but can occupy adult neural territories after transplantation (Figures 5G-5H).

Finally, to evaluate injury responsiveness, recipient colonies received a targeted mechanical injury to the neural gland prior to small-cell transplantation. Confocal imaging at 2 and 9 dpi demonstrate a dramatic accumulation of DiD-labelled small cells at the injured neural gland, adjacent brain-associated regions, along nerves, and in dorsal-organ-associated territories (Figures 5K–5M). Because these experiments were not designed to quantitatively measure directed recruitment, this observation is interpreted as injury-associated accumulation rather than evidence of selective injury-directed homing (Figures 5J-5N).

Together, these transplantation experiments show that neural-complex-derived small cells can persist after vascular delivery and display distinct *in vivo* localization patterns, including association with developing bud territories and localization within neural-complex-associated tissues of the replacement generation. Their localization within bud-associated and neural-complex-associated regions, together with their accumulation around injured neural-complex territories, is consistent with differential tissue association during development and regeneration, while leaving open the precise molecular identity, lineage potential, neuronal differentiation, self-renewal capacity, and long-term functional contribution of the transplanted cells.

## DISCUSSION

Recent single-cell studies across diverse invertebrate and chordate models have revealed fundamental principles governing neuronal identity, highlighting conserved regulatory modules that span cnidarians, annelids, and invertebrate chordates (Bump and Lubeck, 2023; Cao et al., 2019; Dong et al., 2024; Lamanna et al., 2023; Paganos et al., 2021; Pang et al., 2025; Sharma et al., 2019). However, virtually all comparative neurological datasets describe adult nervous systems that are structurally and anatomically static. In contrast the adult central nervous system of *B. schlosseri* operates within an obligate, lifelong cycle of continuous degeneration and *de novo* regeneration.

In this context, *B. schlosseri* provides a complementary chordate model for asking how neural cell states are maintained, remodeled, and reassembled during continuous adult tissue renewal (Anselmi et al., 2022; Kowarsky et al., 2021). This continuous renewal provides a direct counterpoint to vertebrate systems, where adult neurogenesis is severely constrained. In mammals, endogenous NSCs are restricted to specialized niches, principally the subgranular zone of the dentate gyrus and the subventricular zone (Codega et al., 2014; Eriksson et al., 1998; Gage, 2000; van Praag et al., 1999). Within these niches, single-cell and multi-omic profiling reveals that the vast majority of adult NSCs reside in deep, cell-cycle-arrested quiescence (Audesse and Webb, 2020; Codega et al., 2014). Furthermore, prospective isolation of human neural stem and progenitor cell lineages demonstrates that baseline activation and self-renewal capacities decline exponentially following infancy (Liu et al., 2023; Liu et al., 2026). During mammalian aging, neurogenic niches become locked within an inactive, glial-and ependymal-like microenvironment that traps non dividing stem cells in permanent dormancy and curtails endogenous repair (Maimon et al., 2024; Terreros-Roncal et al., 2021). *B. schlosseri* circumvents these bottlenecks by maintaining an accessible, cyclically rebuilt chordate nervous system supported by an active progenitor pool.

Our single-cell transcriptomic atlas and spatial reconstructions demonstrate that the adult *B. schlosseri* neural complex is not a simple, isolated cerebral ganglion, but an integrated, multi-compartment functional territory. Single-cell profiling resolved six transcriptionally distinct cell states encompassing neuroepithelial progenitors (*SOX2*/*MSI1*/*NOTCH1*), differentiated cholinergic and excitability-defined neurons (*NALCN*/*CHAT*), regulatory neuroendocrine-like cells (*GHSR*/*GATA2*), metabolic neural gland cells (*CYP2C5*/*SFTPA1*), innate immune effectors (*CLEC4C*/*CALHM6*) and transitional patterning compartments (*HOXA4*/*NOS1*). Multiplexed HCR-FISH spatially validated these programs across the ganglion cortex, neural gland, ciliated duct, dorsal organ, and peripheral sensory epithelia, proving that transcriptional heterogeneity directly mirrors anatomical architecture. Together with the enrichment of *APP*, *NEUROG1*, and PH3-positive cells in neural-gland-associated territories, these data suggest that the neural gland and dorsal structures participate in neural-complex remodeling during blastogenesis. Because glial identities in adult ascidians remain unresolved, epithelial-like or support-associated states should be interpreted conservatively as neural-complex-associated non-neuronal populations rather than canonical vertebrate glia. Nevertheless, recent work in *Styela plicata* suggests that support-cell-like programs may exist during tunicate CNS regeneration (de Abreu et al., 2022; Medina et al., 2015).

A central finding of this study is the resolution of an uncollapsed chordate neurogenic continuum that separates an immature progenitor pool from committed precursors. Cluster 0 combines markers associated with neural immaturity, progenitor maintenance, epithelial organization, and cell–cell adhesion, including *SOX2*, *MSI1*, *NOTCH*/*HES* pathway components, and *CLDN* family members. *SOX2* is essential for maintaining multipotency and preventing premature differentiation in neural stem cells (Arnold et al., 2011; Martello and Smith, 2014; Mercurio et al., 2019; Zhang and Cui, 2014), while Musashi-1 (*MSI1*) post-transcriptionally maintains the undifferentiated state (Okano et al., 2005; Sakakibara et al., 1996) and *NOTCH*-*HES* signaling sustaining neural stem cells and preventing precocious neurogenesis, both during development and in the adult brain (Imayoshi et al., 2010; Kageyama et al., 2008; Lampada and Taylor, 2023; Louvi and Artavanis-Tsakonas, 2006; Zhou et al., 2022). Where detected, *VIM* further supports this assignment, as vimentin is linked to neuroepithelial and radial glia-like progenitor identities, marking immature neural states with structural and regulatory roles in developing nervous systems (Chen et al., 2018); radial glia expressing such intermediate-filament programs are now recognized as the principal neural stem cells of the vertebrate CNS (Kriegstein and Alvarez-Buylla, 2009). Together, these markers suggest that part of the neural-complex-associated population is organized around a conserved progenitor-like and neuroepithelial regulatory logic rather than representing a nonspecific pool of undifferentiated cells. This interpretation is further supported by previous work showing that *SOXB1*/*SOX2*-like programs in *B. schlosseri* are linked to budding tissues, stemness-associated regulation, and the broader chordate framework of Yamanaka-like factors (Vanni et al., 2022).

In contrast, cluster 2 combines mature cholinergic markers (*CHAT*, *ACHE*) and membrane excitability-associated genes (*NALCN*, *UNC79*, *UNC80*) with the basic helix-loop-helix transcription factor *NEUROG1*. Across bilaterians and solitary tunicates such as *Ciona*, neurogenins act as master proneural determination factors that drive cell-cycle exit and neuronal commitment (Bertrand et al., 2003; Stolfi and Levine, 2011). Cell-cycle and trajectory analyses suggest that progenitor-like and neuronal states in the adult neural complex are not static, discrete populations, but components of a dynamic neurogenic program. The coexistence of cycling and non-cycling *SOX2*-positive cells is consistent with a heterogeneous progenitor compartment, whereas the strong proliferative signature of *NEUROG1*-positive cells suggests that neuronal commitment in *B. schlosseri* may remain closely coupled to cell-cycle activity. The transcriptional connection between clusters 0 and 2 further supports a model in which progenitor-like and proneural/neuronal states represent sequentially related stages of neural renewal rather than independently maintained cell identities.

The less-defined cluster 1 may represent a regulatory or interface state, whereas clusters 3, 4, and 5 likely capture components of the broader neural-complex-associated microenvironment, including epithelial/glandular, immune-like, and transitional programs. The immune/hemocyte-like cluster may be particularly relevant during takeover, when neural tissue remodeling and clearance occur, although a contribution from resident, infiltrating, or dissection-associated hemocytes cannot be distinguished at this stage.

Crucially, our spatial mapping and mitotic profiling revealed that stem and progenitor cells are not randomly distributed throughout the ganglion medulla, but are compartmentalized within specific anatomical structures and brain-adjacent microenvironments. In the adult zooid, HCR-FISH mapped *SOX2* to discrete, isolated mantle cells immediately surrounding the cerebral ganglion cortex, suggesting a spatially restricted perineural progenitor-like compartment. Furthermore, active mitotic proliferation identified by PH3 staining in adult zooids was strictly restricted to early blastogenetic stages (Days 1–3) and localized to the dorsal organ and the ciliated duct/neural gland region. In the developing budding generations that replace the adult brain, *NEUROG1* prominently marks the dorsal tube of primary buds, the transient neurogenic primordium that delaminates to form both the cerebral ganglion and the neural gland *de novo* (Burighel et al., 1998; Prünster et al., 2019). Finally, secondary buds serve as the primary colonial target zones where circulating progenitors accumulate to seed nascent tissues (Levy et al., 2025; Voskoboynik et al., 2008).

Our *in vivo* transplantation experiments suggest that FACS-isolated neural-complex small cells possess intrinsic migratory and homing competencies: they selectively migrate through the colonial circulation into secondary buds, stably incorporate into the cortex, medulla, and nerve roots of the *de novo* regenerated cerebral ganglion of the replacement generation, and accumulate robustly at sites of neural gland injury. Together with previous work on endostyle, germline, cell-island, and ampullar niches, these data place the neural complex within a broader colony-wide logic of trafficking and niche occupancy (Levy et al., 2025; Rinkevich et al., 2013; Rosental et al., 2018; Vanni et al., 2023; Voskoboynik et al., 2008). These findings therefore support the presence of neural-complex-derived cells with *in vivo* persistence and tissue-localization capacity, while future lineage-resolved approaches will be required to determine whether these cells represent the progenitor-like states identified transcriptionally and whether they contribute directly to neural differentiation.

Because *B. schlosseri* completely resorbs its adult zooids weekly, all differentiated somatic tissues, including every mature neuron, synapse, and sensory receptor, are eliminated during each blastogenetic takeover (Anselmi et al., 2022; Kowarsky et al., 2021; Thompson et al., 2022). Consequently, in the chronologically aged colonies analyzed in our study (cultured continuously in mariculture for 10 to nearly 20 years, spanning over 500 to 1,000 successive blastogenetic generations), every single adult brain analyzed is the direct product of massive, cumulative stem cell-mediated turnover. This biological reality fundamentally reframes our understanding of chordate brain aging. In traditional mammalian models, post-mitotic neuronal aging reflects accumulated intracellular damage within non-renewed cells (López-Otín et al., 2013). In *B. schlosseri*, because adult neurons are never more than seven days old, age-related neurological decline, including progressive neuron loss, siphon reflex deficits, and altered gene expression (Anselmi et al., 2022; Anselmi et al., 2023; Bocci et al., 2024), must be intrinsically driven by the aging and clonal exhaustion of the ancestral stem cell pool that repeatedly generates them. This mirrors mammalian hematopoietic aging, where lifelong clonal expansion of altered stem cells impairs downstream lineage potential (Beerman et al., 2010; Pang et al., 2011).

Bulk temporal transcriptomics revealed that young adult brains follow a tightly coordinated blastogenetic trajectory, where excitability machinery (*NALCN*, *UNC79*, *UNC80*, *CHAT*) peaks during mid-cycle to support behavioral responsiveness, while tight-junction remodeling (*CLDN1*, *CLDN7*) and immune effectors (cluster 4 hemocytes) expand at Day 7 during takeover. Automated Deconvolution Augmentation of Profiles for Tissue Specific cells (ADAPTS) deconvolution revealed that in aged colonies (spanning up to 6,845 days in mariculture), this developmental coordination becomes desynchronized, exhibiting altered proportional representation of progenitor (*SOX2*), neuronal (*NALCN*), and immune (*CLEC4D*) programs during early blastogenesis. Concurrently, *APP* exhibited a continuous, monotonic upregulation from Day 1 to Day 7, mapping strongly to the neural gland and regressing tissues. While mammalian APP is centrally studied in Alzheimer’s disease neuropathology, APP executes physiological roles in synaptic remodeling and cell stress responses (Müller et al., 2017). In *B. schlosseri*, cyclical *APP* elevation accompanies non-pathological tissue resorption and *de novo* budding, suggesting that amyloid cascades originally evolved as conserved modules regulating physiological cell clearance and neural remodeling.

In conclusion, this work establishes that lifelong CNS regeneration in *B. schlosseri* is sustained by conserved progenitor-like states operating within defined anatomical niches and dynamically linked to proneural and neuronal programs during successive rounds of CNS renewal. Future investigations must elucidate the specific chemotactic receptors (such as conserved *SDF-1*/*CXCR4* or integrin axes; (Wright et al., 2001)) that direct progenitor homing to secondary buds, uncover the mechanisms that preserve chordate stem cell lineages across thousands of asexual cycles (Domen et al., 2026) and test whether bioelectrical stimulation or systemic circulatory interventions can reverse the epigenetic desynchronization of the aging neural stem cell pool (Domen et al., 2026). Resolving these mechanisms in *B. schlosseri* will deepen our understanding of chordate neurogenesis and provide fundamental evolutionary strategies to overcome the dormancy barriers that restrict neural repair in the aging human brain.

## MATERIALS AND METHODS

### Animal husbandry and staging

Colonies of *B. schlosseri* were maintained under standard mariculture conditions and staged according to the blastogenic cycle. To define the cellular composition of the adult *B. schlosseri* CNS (neural complex), we isolated brains from adult zooids during early blastogenesis (day 1 and day 2). This specific developmental stage was selected because total neuronal numbers increase significantly during this early adult phase (Anselmi et al 2022), making it the most informative stage in which to capture active progenitor-like and proliferative cellular states.

### Neural complex dissection and cell dissociation

The neural complex, including the cerebral ganglion and associated neural gland region, was manually dissected under a stereomicroscope from live animals and processed immediately for cell dissociation. A total of 50 adult neural complexes (CNS samples) were dissected, all derived from colonies of the same genotype (6941c) and collected at 221 days of age. Freshly dissected adult brains of *B. schlosseri* were dissociated enzymatically using Accumax (Innovative Cell Technologies), a cell detachment solution with proteolytic and collagenolytic activities. Dissociation conditions had been optimized at 24°C, and adult brains were incubated for 1 h. During dissociation, samples were intermittently triturated by pipetting to facilitate disruption of cell aggregates and residual tunic material. Following enzymatic treatment, cells were resuspended in salinity adjusted medium prepared by mixing 10x HBSS and 1x Medium 199 at a 1:2.03 volume ration, and filtered through a 40 _μ_m nylon cell strainer to obtain a clean single-cell suspension for FACS and downstream single-cell RNA-seq.

### Fluorescence-activated cell sorting and index sorting

Live cells derived from dissociated neural complexes were subjected to FACS with an indexed single-cell sorting workflow. Based on forward-and side-scatter parameters, brain-enriched cell populations were separated into two major gates corresponding to relatively small and relatively large cells (Fig 1f). Individual cells from these gates were sorted into 96-well plates for downstream single-cell RNA-seq. The sorting strategy was designed not only to isolate single viable cells for sequencing, but also to retain index information, allowing each sequenced cell to be mapped back to its original FACS position and size gate. This design was later used to relate transcriptional identities to the initial physical sorting parameters and to visualize the distribution of “small” and “large” gates across the UMAP embedding.

### Smart-seq3 library preparation and sequencing

Single cells obtained by index sorting were processed using Smart-seq3 for full-length transcriptome profiling with UMI-aware quantification. The resulting libraries were sequenced as paired-end reads and used to generate digital gene expression matrices for downstream quality control and clustering.

Index-sorted single cells were processed using a modified Smart-seq3 workflow for full-length transcriptome profiling with UMI-aware quantification as described in Levy et al. 2026. Individual cells were sorted with a Sony MA900 FACS into 96-well plates containing 2 _μ_l of lysis buffer. Immediately after sorting, plates were centrifuged at 3000 × g for 30 s, snap-frozen on dry ice, and stored at −80°C until processing. For reverse transcription, frozen lysis plates were thawed on ice and heated to 72°C for 3 min to promote annealing of the oligo-dT30VN primer, followed by immediate cooling on ice. Reverse-transcription mix was then added to each well using a Mantis liquid handler (Formulatrix), and reactions were incubated in a C1000 Touch Thermal Cycler (Bio-Rad) at 42°C for 90 min, followed by 70°C for 15 min to terminate the reaction. cDNA amplification was performed by adding PCR mix to each well and cycling as follows: 98°C for 3 min; 25 cycles of 98°C for 20 s, 67°C for 15 s, and 72°C for 6 min; and a final extension at 72°C for 5 min. Amplified cDNA was purified using 0.65–0.75× calibrated AMPure XP beads (Beckman Coulter) to remove reaction components and fragments smaller than approximately 400 bp, and was eluted in 12.5 _μ_l ultra-pure water. Plate clean-up was performed using a Bravo automated liquid handling platform (Agilent). For quality control, 1 _μ_l of purified cDNA from each well was analyzed for concentration and fragment-size distribution using a Fragment Analyzer (Advanced Analytical). cDNA from 96-well plates was then transferred into 384-well plates using a Mosquito X1 liquid handler (SPT Labtech), and concentrations were normalized with ultrapure water so that each well did not exceed 1 ng/_μ_l. Normalized cDNA was used for Illumina library preparation. Tagmentation reactions were assembled using 0.8 _μ_l cDNA and 1.2 _μ_l Tn5 transposase mix, and were stopped by addition of 0.4 _μ_l neutralization buffer containing 0.1% SDS. Indexing PCR was performed using custom i5 and i7 dual-index primers together with KAPA HiFi HotStart ReadyMix (Kapa Biosystems). PCR cycling consisted of 72°C for 3 min, 95°C for 30 s, followed by 11 cycles of 98°C for 10 s, 67°C for 30 s, and 72°C for 1 min. Libraries from each 384-well plate were pooled by combining 1 _μ_l from each well and purified using 0.8× AMPure XP beads. Each 384-cell library pool was assessed by Fragment Analyzer, normalized by concentration, and combined into a final pooled library. After a final AMPure XP bead-based purification and concentration step, the pooled library was sequenced on an Illumina NovaSeq 6000 S4 flow cell to generate approximately 1.5 million paired-end 2 × 150 bp reads per cell.

### Read processing, QC, clustering and annotation

The raw dataset comprised 676 sequenced cells. Quality control filtering was then applied to remove low-quality cells and likely doublets, retaining 470 high-quality cells for downstream analysis. Cell filtering was based on the number of detected genes and total transcript counts per cell, using fixed lower thresholds of 1000 counts and 100 genes together with upper cutoffs defined by median + 5 × median absolute deviation (MAD). After cell-level filtering, genes with detectable expression in at least one retained cell were kept for downstream analysis. This reduced the feature space from 44,727 genes to 17,118 genes. The retained dataset therefore represented 69.5% of the original cells and 38.3% of the original gene set.

Filtered data were analyzed in Seurat. Counts were normalized using Seurat’s global-scaling normalization approach (NormalizeData), in which gene counts for each cell were divided by total cellular counts, multiplied by a scale factor of 10,000, and log-transformed. Highly variable genes were identified using FindVariableFeatures, and data were scaled to zero mean and unit variance prior to dimensionality reduction. Intrinsic dimensionality analysis suggested that approximately 4–5 dimensions captured the main biological structure of the dataset, and clustering was therefore performed using the Louvain algorithm at resolution 0.2 on the first five principal components, yielding a silhouette score of 0.44 and six transcriptionally distinct clusters. For visualization, both UMAP and t-SNE embeddings were generated from the selected PCs.

Indexed FACS information was retained for each sequenced cell, allowing the physical properties recorded during sorting to be linked to the final transcriptomic identity of the same cell. In particular, the original small-and large-cell FACS gates were projected onto the UMAP embedding to examine whether size-based sorting corresponded to distinct transcriptional populations.

Unsupervised clustering resolved six transcriptionally distinct populations in the *B. schlosseri* neural complex. Cluster-enriched markers were identified using Seurat’s FindAllMarkers function with a Wilcoxon rank-sum test, retaining positive markers detected in at least 25% of cells in the target cluster, showing at least two-fold enrichment, and having a Bonferroni-adjusted *p* value < 0.05. Because many transcripts in the *B. schlosseri* genome remain incompletely annotated, final cluster annotation was not based solely on top-ranked differentially expressed genes. Instead, cluster identities were assigned by integrating data-driven marker lists, curated conserved “anchor” genes representing known cell-state programs, and ontology/enrichment-based interpretation of annotated cluster-enriched transcripts.

This annotation strategy was used to define the final cluster nomenclature, emphasizing a neuroepithelial/progenitor-like program, a *GHSR*/*GATA2*-like receptor/transcription-factor-enriched state, a neuronal/cholinergic and excitability-rich population, a secretory/metabolic epithelial-like population, an immune/hemocyte-like population, and a transitional transport/patterning-enriched state. Extended marker visualizations and the annotation framework were used to support the final cluster identities.

To interpret the atlas in a comparative neurobiological framework, the expression of selected *B. schlosseri* homologs associated with vertebrate neurogenesis, neurotransmission, excitability, and neurodegeneration was projected onto the UMAP embedding. These included genes such as *NEUROG1*, *SOX2*, *CHAT*, *NALCN*, *UNC79*, *UNC80*, *CNTNAP2*, *CDH23*, *TRPC3*, *COMT*, *TSPAN6*, *UMOD*, and *APP*.

### Gene annotation and cross-species nomenclature

Gene identities were assigned according to the existing *B. schlosseri* genome annotation (Voskoboynik et al., 2013), in which predicted gene models were associated with putative homologs from annotated vertebrate gene sets based on sequence similarity. Because *B. schlosseri* is a non-model organism with incomplete functional annotation and lineage-specific gene duplications, these assignments were used as putative functional annotations rather than as evidence of strict one-to-one orthology.

Throughout the manuscript, conserved gene symbols (e.g., *SOX2*, *CHAT*, *NALCN*) therefore refer to *B. schlosseri* gene models annotated as homologous to the corresponding vertebrate genes. Gene annotations were used to support cluster interpretation and comparison with conserved developmental and neural programs. Such cross-species comparisons were considered hypothesis-generating and were not used as evidence of conserved gene function in the absence of experimental validation.

### HCR probe design and HCR-FISH

HCR-FISH target genes were selected from the scRNA-seq marker analysis to represent the major transcriptional states identified in the adult neural-complex atlas. Because candidate genes differed in whether they were strongly expressed in rare subsets or more broadly detected across clusters, target selection was guided by both average expression and percent-positive metrics. This approach allowed us to prioritize genes representing both strongly enriched, spatially restricted populations and more broadly distributed cluster-associated programs (Table S4).

Antisense DNA probe sets specific for selected target genes were designed against coding regions identified through the in silico pipeline. Probe sets associated with initiator B1 were ordered as lyophilized oligonucleotides from Integrated DNA Technologies. The complete probe sequences are provided in Table S5. *APP* and *NEUROG1* probe sets were designed and synthesized directly by Molecular Instruments. Upon arrival, probes were resuspended in nuclease-free water to a final concentration of 0.5 _μ_M and stored at −20°C until use. After acclimation and once colonies reached the desired developmental stage, one system was excised from each selected colony and fixed overnight at 4°C in 4% paraformaldehyde (PFA) in 1× PBS. Samples were then washed three times in 1× PBS for 10 min each, dehydrated through a graded methanol series in 1× PBS, and stored in 100% methanol at −20°C until use.

Methanol-stored samples were rehydrated through decreasing methanol concentrations in 1× PBS and washed first in 1× PBS for 10 min and then in 0.1% Tween-20 in PBS (PBT) for 30 min. Samples were post-fixed in 4% PFA in 1× PBS for 25 min and washed five times in PBT for 5 min each. Samples were pre-incubated in hybridization buffer (Molecular Instruments) for 30 min at 37°C and then incubated overnight at 37°C in hybridization buffer containing antisense probe. Probe concentration was 0.002 _μ_M for the *APP* probe, used as a positive control, and 0.02 _μ_M for all other probes. Negative controls were incubated in hybridization buffer without probe. The following day, excess probe was removed by four 15 min washes in probe wash buffer (Molecular Instruments). Samples were then incubated in amplification buffer (Molecular Instruments) for 30 min and subsequently transferred to hairpin solution prepared by mixing 2 _μ_L of hairpin 1 and 2 _μ_L of hairpin 2 in 100 _μ_L of amplification buffer. Amplification proceeded overnight at room temperature in the dark. Because HCR hairpins are highly photosensitive, all post-amplification steps were carried out under reduced light conditions. After amplification, samples were washed five times in 0.1% Tween-20 in 5× SSC (SSCT) for 5, 5, 30, 30, and 5 min, respectively. Nuclei were counterstained with Hoechst 33342 in 1× PBS at a final concentration of 2 _μ_g/_μ_L for 3 h. Excess dye was removed by two 10 min washes in 1× PBS. Samples were then incubated in 30% glycerol for 30 min followed by 70% glycerol for 30 min and mounted on microscopy slides with the oral siphon oriented upward for confocal imaging.

Negative controls lacking probe were used to assess tissue autofluorescence and nonspecific background. The APP probe was used as a positive control because previous data had shown robust and reproducible expression in the neural gland region throughout the blastogenetic cycle, allowing validation of protocol performance.

### Cell-cycle scoring

Cell-cycle state was computationally inferred from the single-cell transcriptomes using the CellCycleScoring function implemented in Seurat, following approaches previously applied to single-cell datasets in invertebrate systems (Brunet Avalos et al.; Gordon et al., 2025). Reference gene sets associated with S phase and G2/M phase provided with Seurat were matched to the annotated *B. schlosseri* features, including transcript variants sharing the corresponding gene names. This resulted in 31 S-phase-associated and 26 G2/M-associated features represented in the dataset. Seurat was then used to calculate S and G2/M expression scores for each cell and to assign a predicted cell-cycle phase (G1, S, or G2/M). The proportion of cells assigned to each phase was calculated for individual transcriptional clusters, and the proliferative fraction was defined descriptively as the proportion of cells assigned to S or G2/M.

To examine cell-cycle state in relation to progenitor-and proneural-associated markers, cells were additionally classified according to *SOX2* and *NEUROG1* expression. A cell was considered positive when normalized expression greater than zero was detected for at least one matched *SOX2* or *NEUROG1* feature. Cell-cycle phase distributions were then compared among *SOX2*-positive, *NEUROG1*-positive, double-positive, and negative cells. Because this approach infers cell-cycle state from transcriptional signatures rather than directly measuring DNA content or mitotic activity, phase assignments were interpreted as predicted cell-cycle states. Active mitosis was independently assessed by PH3 immunostaining as described below.

### Trajectory inference and pseudotime analysis

Trajectory inference was performed using Slingshot on the final Seurat object containing 470 cells. The first five principal components, corresponding to the dimensional space used for clustering, were used for trajectory inference. An initial unrooted analysis was performed without specifying a starting cluster to assess relationships among the six transcriptional clusters independently of their biological annotation. This analysis identified a direct connection between progenitor-like cluster 0 and neuronal cluster 2. A second analysis was then performed using cluster 0 as a biologically informed starting state, based on its enrichment for *SOX2*, *MSI1*, *HES*, and *NOTCH*-associated genes. Pseudotime values were extracted for the lineage connecting clusters 0 and 2; the resulting lineage contained 149/154 cluster 0 cells and 75/75 cluster 2 cells. Pseudotime was inferred in five-dimensional PCA space and projected onto the existing UMAP coordinates for visualization only. To examine transcriptional changes along the inferred 0–2 axis, lineage-associated cells were divided into ten equal-frequency pseudotime bins, and mean normalized expression was calculated within each bin for selected progenitor-associated and proneural/neuronal genes. Expression values were subsequently scaled independently for each gene across pseudotime bins for visualization. To assess within-cluster changes in *SOX2* and *NEUROG1* detection, cluster 0 and cluster 2 cells, respectively, were divided into pseudotime quartiles. Cells were classified as positive when normalized expression was greater than zero. Differences in the proportion of positive cells across quartiles were tested using Fisher’s exact test, followed by pairwise proportion tests with Benjamini–Hochberg correction for multiple comparisons.

### PH3 Immunostaining

PH3 is a widely used mitotic marker, including in tunicates, and therefore provides a spatial readout of active mitosis rather than lineage identity (Hendzel et al., 1997; Goto et al., 1999; Schulmeister et al., 2007; Gasparini et al., 2013; Kawamura et al., 2015). 12 colonies at defined blastogenetic stages were fixed overnight at 4°C in 4% paraformaldehyde (PFA) in 1× PBS. Specifically, three colonies at day 1, two colonies at day 2, three colonies at day 3, one colony at day 4, two colonies at day 5, and one colony at take-over (TO) were processed. After fixation, samples were washed in PBS, permeabilized, and blocked in 1% BSA in PBS-T. Samples were then incubated with primary antibodies against phospho-Histone H3 (rabbit anti-phospho-histone H3 Ser10, 1:200) and _α_-tubulin (mouse anti-_α_-tubulin, 1:1000), followed by species-appropriate fluorescent secondary antibodies. Nuclei were counterstained with Hoechst or DAPI. After mounting in glycerol, samples were imaged by confocal microscopy and examined for the presence and localization of PH3-positive mitotic cells within the adult, primary bud, and secondary bud neural complexes across developmental stages.

### Confocal imaging and image processing

Whole-mount samples were imaged to assess signal across the entire colony rather than only in the neural complex. For each sample, all adult zooids and buds present in the system were examined. Images were acquired using a Zeiss LSM 900 Airyscan 2 confocal microscope at the University of Padova Imaging Facility. For some samples, Z-stacks were collected with an average of 25 optical sections at 1 _μ_m intervals. Images were processed using ImageJ (version 1.54p).

### Bulk brain collection and RNA-seq dataset generation

Bulk neural-complex samples were collected from *B. schlosseri* colonies spanning defined blastogenetic stages and chronological ages (See Table S3). Neural complexes were manually dissected from individual adult zooids using insulin syringes, following the general tissue-collection procedure described previously (Anselmi et al. PNAS). For each bulk RNA-seq sample, 3 or 4 neural complexes from each colony were pooled before RNA extraction. Samples were homogenized using a mechanized Kontes tissue grinder and pestle in the presence of lysis buffer from the RNeasy Micro Kit. Total RNA was extracted according to the manufacturer’s protocol and subsequently cleaned and concentrated using the RNA Clean & Concentrator-5 kit. RNA quantity and integrity were assessed using an Agilent 2100 Bioanalyzer, and samples meeting the required RNA-quality criteria were selected for library preparation.

Raw sequencing reads were processed. Reads were aligned to the *B. schlosseri* reference genome (Voskoboynik et al 2013) and last annotation version (Levy et al 2025), and gene-level counts were generated using. The resulting raw count matrix was used for the downstream quality-control, differential-expression, stage-associated, and single-cell-reference integration analyses described below.

### Bulk RNA-seq dataset curation and analysis

Bulk RNA-seq data from *B. schlosseri* brains were analyzed to assess transcriptional variation across blastogenetic stages and colony age. Samples were classified into two biological groups based on developmental stage and colony age: young adult brains, old adult brains. After quality filtering, the bulk brain RNA-seq dataset used for downstream analyses comprised young adult samples distributed across stage A (n = 8; 77, 82, and 188 days), stage B (n = 10; 47, 77, 131, and 139 days), stage C (n = 11; 66, 82, 189, and 243 days), and stage D (n = 5; 72 and 82 days), as well as old adult samples at stage A (n = 4; 6845 days) and stage C (n = 2; 3721 days). Young adult samples were further subdivided into stages A, B, C, and D according to the blastogenetic cycle, whereas old adult samples were retained for age-matched comparisons at stages A and C.

Sample-level quality control was performed on all brain bulk RNA-seq samples using library size, number of detected genes, principal component analysis (PCA), and mean sample-to-sample Pearson correlation. For each sample, we quantified total library size, total number of detected genes, and mean correlation with the remaining samples after log-transformation of count data. Samples with extremely low library size, very low gene detection, and/or globally poor correlation relative to the dataset were considered technical outliers and excluded from downstream analyses. After filtering, the cleaned dataset comprised young adult samples at stages A–D, old adult samples at stages A and C. This cleaned dataset was used for all downstream age-and stage-associated analyses.

For exploratory analyses, raw counts were transformed as log2 (count + 1). PCA and sample correlation heatmaps were generated on filtered count matrices to assess the global structure of the dataset. In young adult samples, exploratory analyses were interpreted as describing stage-associated transcriptional patterns, because genotype was partially nested within the blastogenetic stage. Age-matched comparisons (young A vs old A; young C vs old C) were interpreted as cleaner tests of age-associated variation within the same stage. Differential expression analyses were performed in DESeq2 using rounded raw count matrices. For age-matched two-group comparisons, including old A vs young A, old C vs young C, and old C vs old A, DESeq2 models used a simple design with the relevant grouping variable. Wald statistics were used for pairwise contrasts, and results were ranked by adjusted *p* value. Genes with positive log2 fold change were interpreted as more highly expressed in the first group of the contrast, whereas negative values indicated higher expression in the reference group.

To identify genes associated with blastogenetic stage across young adult brains, we used a likelihood ratio test (LRT) with the design ∼ stage4 and reduced model ∼ 1. This analysis identified genes whose expression varied significantly across stages A–D without restricting the test to any single pairwise contrast. These genes are referred to throughout the manuscript as stage-associated bulk DE genes.

To interpret bulk brain transcriptional changes in a cell-state-informed framework, we used marker genes derived from the single-cell RNA-seq atlas of the adult neural complex. The scRNA-seq dataset had been generated from 676 Smart-seq3-profiled cells, filtered to 470 high-quality cells, normalized and clustered into six transcriptionally distinct populations, annotated as neuroepithelial/progenitor-like, *GHSR*/*GATA2*-like, neuronal, secretory epithelial, immune/hemocyte-like, and transitional/patterning-related states. Cluster annotation was based on curated anchor markers and differential expression analysis.

For bulk integration, we defined compact marker sets for each of the six scRNA-derived clusters and matched them to the curated bulk annotation. For each bulk sample, a cluster signature score was calculated as the mean log2-transformed expression of the marker genes associated with each scRNA-defined cluster. These signature scores were then compared across young stages A–D and across aging-related contrasts (young A vs old A, young C vs old C, and old A vs old C). Nonparametric Kruskal–Wallis tests were used for comparisons across young stages, and Wilcoxon rank-sum tests were used for pairwise aging-related signature comparisons.

In parallel, stage-associated bulk DE genes identified in the young dataset were intersected with scRNA-derived cluster marker sets to determine which cell-state programs contributed most strongly to stage-associated bulk remodeling. Compact heatmaps were generated using representative top stage-associated DE genes grouped by scRNA-defined cluster assignment.

### Cell isolation, transplantation and imaging

For live *in vivo* tracking, small-cell fractions were analyzed in six recipient colonies/subclones, whereas large-cell fractions were analyzed in three recipient colonies/subclones. Recipient colonies were imaged at 3 and/or 5 days post injection (dpi), and DiD-labelled cells were scored across adult cell islands, primary buds and secondary buds when present.

For confocal imaging, non-injured small-cell transplantations were analyzed in three recipient colonies/subclones at 2–7 dpi. Injury experiments were performed after transplantation of DiD-labelled small-cell fractions and included neural gland damage or brain damage. These injury experiments were analyzed in three identified recipient subclones at 2, 7, or 9 dpi. The number of imaged fields and positive fields for each condition, time point, and anatomical compartment is reported in Table S3.

Brain-enriched cell suspensions were obtained from dissected donor animals at stage A by mechanical dissociation, as described above. Cells were sorted using the same gating strategy adopted for the single-cell experiments, separating small-and big-cell populations. After sorting, cells were centrifuged, resuspended in 500 _μ_l of *B. schlosseri* medium, and labeled with 5 _μ_l of Vybrant DiD Cell-Labeling Solution (Invitrogen). Labeled cells were incubated for 1 h at room temperature in the dark, washed twice in *B. schlosseri* medium, first supplemented with 2% FCS and then in serum-free medium, and finally resuspended in 30 _μ_l of fresh serum-free medium prior to transplantation. For transplantation, glass needles were prepared using a P-1000 micropipette puller (Sutter Instruments). Needles were loaded with 5 _μ_l of labeled cell suspension, and injections were performed using a manual micromanipulator (World Precision Instruments) under a stereomicroscope. The number of cells recovered by FACS for each experimental pool was recorded and is reported in table S3. When a sorted cell pool was distributed among multiple recipient colonies, the corresponding cell input per recipient was estimated from the number of sorted cells and the number of recipients. Cells were injected into the proximal ampulla or blood vessels of recipient animals at stage A. Donor and recipient colonies were subclones of the same genotype to ensure compatibility between transplanted cells and host tissues. Recipient colonies were monitored *in vivo* at 3 and 5 days post injection using the BZ-9000 Keyence microscope. As controls, stage A subclonal animals of the same genotype were either left untreated or injected with Vybrant DiD solution in the absence of cells. To examine transplanted cells at higher spatial resolution, recipient colonies were fixed and processed for confocal microscopy (Zeiss LSM700) following cell transplantation. More specifically, donor-derived brain-enriched cell suspensions were prepared, sorted, labeled, and injected into compatible recipient colonies as described above. Two days or 9 days after transplantation, colonies were fixed in 4% paraformaldehyde (PFA), stained with DAPI, and imaged by confocal microscopy to assess the localization of labeled donor-derived cells within host tissues.

To test the behavior of transplanted cells in an injured host tissue, a separate set of experiments was performed in which recipient colonies were subjected to neural gland damage before transplantation. Damage was induced mechanically using a tungsten needle, which was used to selectively injure either the neural gland of recipient animals. Labeled donor-derived cells were injected after injury, and recipient colonies were fixed 1 or 2 days later in 4% PFA, stained with DAPI, and analyzed by confocal microscopy. Control conditions included injured recipient colonies that were not injected with cells.

## SUPPLEMENTARY TABLE

**Table S1. Cluster marker genes.** For each scRNA-seq cluster, marker genes were identified using Seurat and ranked by fold change. Only positive markers were retained, requiring detection in at least 25% of cells in the target cluster, adjusted *p* value < 0.05, and at least two-fold enrichment relative to the remaining cells. The compact table reports the 50 top-ranked annotated marker genes for each cluster together with differential expression statistics, including adjusted *p* value, fold change, and fraction of positive cells. The full ranked output is provided as source data in the file cluster_markers_top50_fc.csv.

**Table S2. Cluster nomenclature and annotation rationale.** Final cluster names were assigned by integrating core differentially expressed markers, secondary supporting markers, and functional enrichment results. Ontology/enrichment support refers to representative over-representation terms obtained from Enrichr analyses of cluster-associated marker sets, together with manual interpretation of marker identity and comparative literature. The table documents the logic used to move from preliminary cluster labels to the final nomenclature adopted in the manuscript.

**Table S3. Sample and dataset summary.** Overview of all samples included in the HCR-FISH analyses, immunostaining and transplantation, including colony origin, developmental stage, age, tissue type, experimental application, and number of zooids, brains, or cells analyzed.

**Table S4. Cluster assignment of HCR target genes based on single-cell expression** For each HCR gene (exact transcript variant), was reported the cluster in which the gene shows the highest average expression (*top_cluster_by_avg*; *avg_exp_avgTop*) and the fraction of cells expressing the gene in that cluster (*pct_pos_avgTop*). It was also reported the cluster in which the gene is expressed in the largest fraction of cells(*top_cluster_by_pct*; *pct_pos_pctTop*) together with the corresponding average expression in that cluster (*avg_exp_pctTop*). Average expression and percent-positive values were derived from Seurat DotPlot summaries across seurat_clusters, enabling identification of genes that are either strong but restricted to a subset of cells (high average, low percent) or broadly expressed (high percent, moderate average).

**Table S5:** List of antisense DNA oligonucleotide probes used for HCR-FISH detection of selected *Botryllus schlosseri* transcripts. Probe sets reported in the table were designed against the corresponding target sequences and synthesized by Integrated DNA Technologies (IDT). For *APP* and *NEUROG1*, the probe sets were designed and synthesized directly by Molecular Instruments; therefore, the individual oligonucleotide probe sequences are not reported. Instead, the target transcript sequence provided to Molecular Instruments and used for probe-set design is included.

## AUTHOR CONTRIBUTIONS

C.A., A.V., and I.L.W. conceived the study. C.A., T.L., A.V., and G.M., contributed to the overall experimental design. C.A. and K.J.I. performed neural-complex dissections. C.A. performed cell dissociation. C.A., T.L., and R.S. performed flow-cytometry sorting. C.A. performed HCR-FISH, PH3 immunostaining, and confocal microscopy. K.J.I. and K.J.P. maintained *Botryllus schlosseri* colonies and contributed to mariculture, staging, and animal preparation. C.A. and T.L. performed single-cell isolation and Smart-seq3 library preparation. A.M.D., N.F.N., and D.L. supported single-cell sequencing and data generation. C.A., J.C.Y, and C.L. performed sequencing analysis, genome/gene annotation integration, and bioinformatic analyses. K.J.P., K.J.I., and C.A. prepared the bulk RNA-seq plates. C.A. and J.C.Y performed bulk RNA-seq integration and visualization with input from Y.V. C.A. performed transplantation experiments and live/imaging-based analysis. C.A., A.V., and G.M. wrote the manuscript. All authors reviewed, edited, and approved the final manuscript.

## ACKNOWLEDGEMENTS

We thank Marialia Coco, Irene Esegio, Francesca Pia Ciffelli, Alisa Zgonnikova and Leyla Yilmaz for their assistance with HCR-FISH experiments, and Elisa Boldrini for her assistance with immunostaining experiments. We thank Stuart Thompson, Laurent Formery, Paul Bump, Tal Raveh, Anna Eastman and Lucia Manni for helpful and valuable discussions. We are grateful to Christopher Lowe for valuable discussions and for providing resources and support that enabled the execution of this project. We also thank Linda Quinn and Tejaswitha Naik for laboratory management and support, and the Chan Zuckerberg Biohub San Francisco for sequencing expertise and support

## SUPPLEMENTARY FIGURES

**Figure S0.**
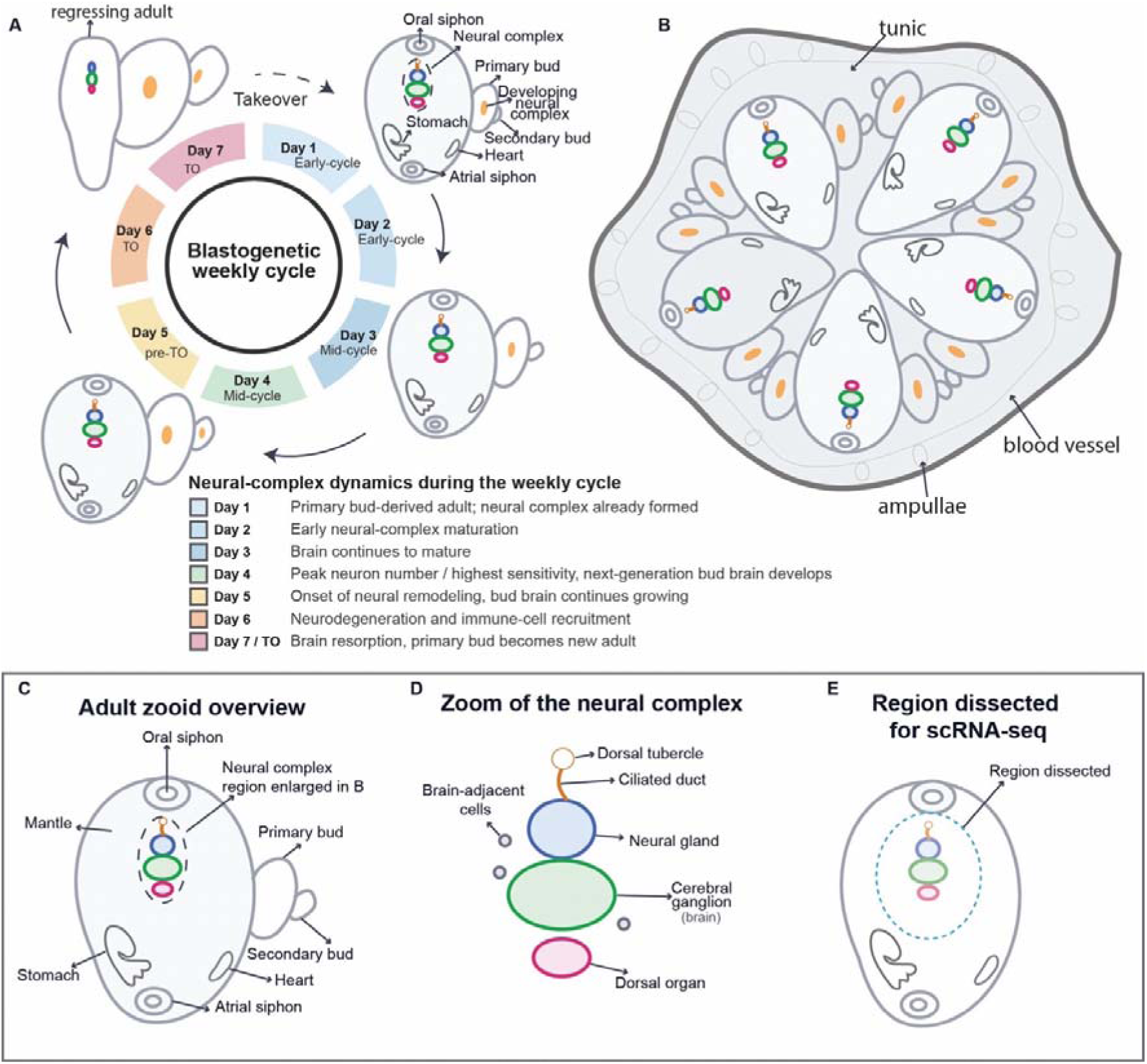
Colony organization, adult zooid anatomy, neural-complex structure, and day-based staging across the weekly blastogenetic cycle of *Botryllus schlosseri*. (A) *Botryllus schlosseri* reproduces sexually through embryogenesis and asexually through blastogenesis. This schematic illustrates the weekly asexual budding cycle, during which clonally related zooids are generated, mature, regress, and are replaced by the next generation. The dashed arrow between Day 7/takeover and Day 1 indicates a generational transition rather than the continuation of the same adult zooid: at takeover, the regressing adult zooid is resorbed, while the primary bud of the previous cycle becomes the new adult zooid. Thus, the Day 1 neural complex corresponds to the brain that developed in the primary bud during the preceding cycle. During Days 1–3, the adult neural complex continues to mature. Around Day 4, the adult brain reaches maximal neuron number and the zooid shows the highest behavioral sensitivity; at the same time, the neural complex of the next generation is developing in the bud lineage. During Days 5–6, the adult brain enters a remodeling and degeneration phase, characterized by decreasing neuron number, increased apoptotic/neurodegenerative signatures, and increased immunocyte association with the neural complex. At Day 7/takeover, the old adult zooid and its brain are resorbed, while the bud generation replaces it. These stage-dependent neural changes were previously characterized in *B. schlosseri*. (B) Stylized view of a zooid within the colonial context. The colony is attached to a substrate, covered by a common tunic, and composed of genetically identical zooids produced by asexual budding. Zooids are connected by a shared extracorporeal blood vasculature ending in peripheral ampullae. This transparent and experimentally accessible vascular network allows live observation and can be used for injection-based manipulation. (C) Overview of an adult zooid showing the oral siphon, atrial siphon, mantle, stomach, heart, primary bud, secondary bud, and the neural-complex region. (D) Enlarged schematic of the neural complex, including the cerebral ganglion/brain, neural gland, ciliated duct, dorsal tubercle, dorsal organ, and associated brain-adjacent cells. (E) Region dissected for Smart-seq3 single-cell RNA-seq, encompassing the neural complex and closely associated tissues.

**Figure S1.**
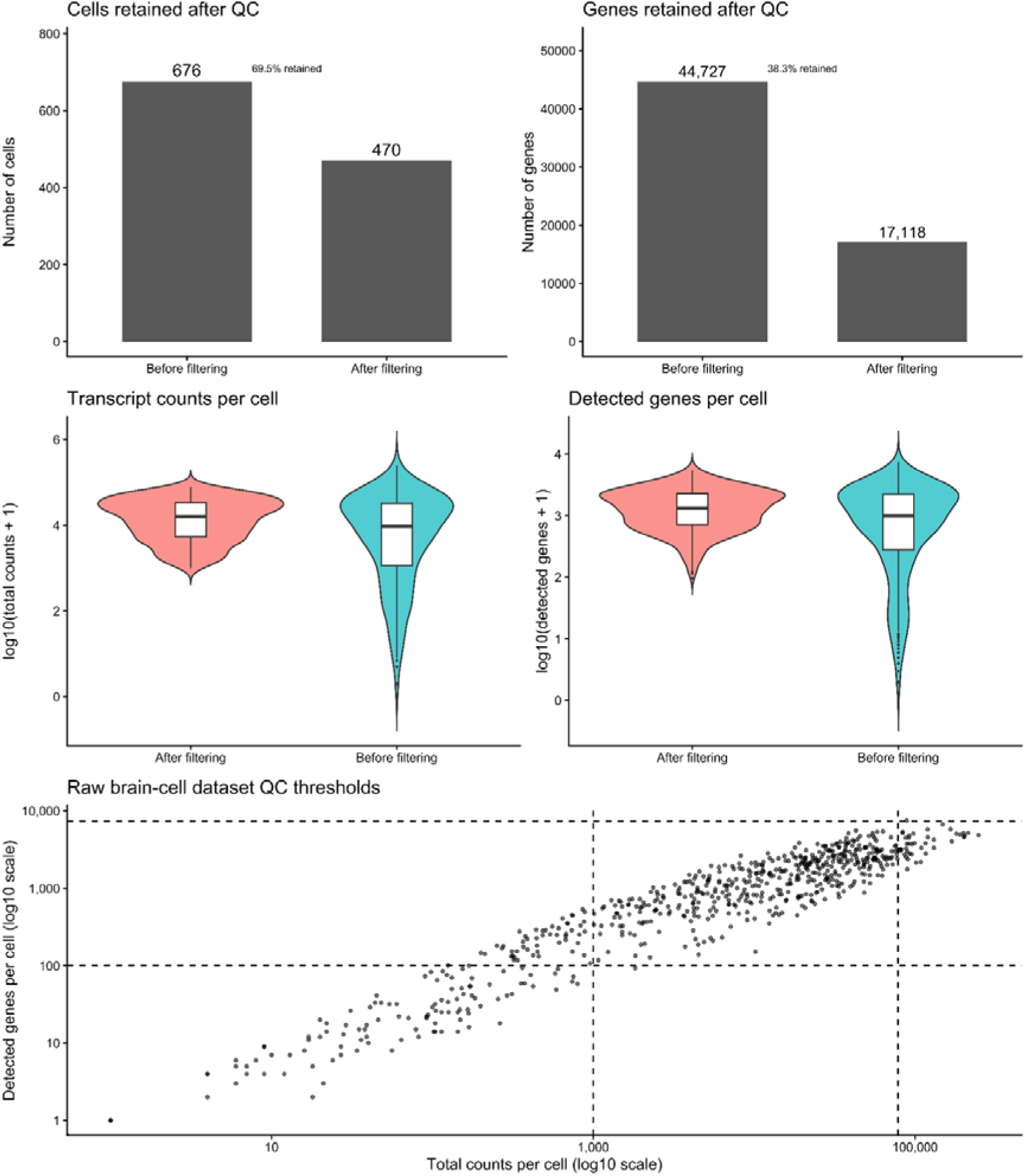
Quality control and filtering of the scRNA-seq dataset. Summary of the quality-control workflow applied to the Smart-seq3 dataset generated from dissected adult *Botryllus schlosseri* neural complexes. The top panels show the numbers of cells and genes before and after filtering, indicating retention of 470 of 676 cells and 17,118 of 44,727 genes. The middle panels show the distributions of total transcript counts and detected genes per cell before and after filtering. The bottom panel shows the raw brain-cell dataset in count–gene space, with dashed lines indicating the thresholds used for filtering. Cells were retained using fixed lower cutoffs together with upper thresholds defined from the raw dataset using a median + MAD-based approach. Exact threshold values and summary statistics are reported in the Table.

**Figure S2.**
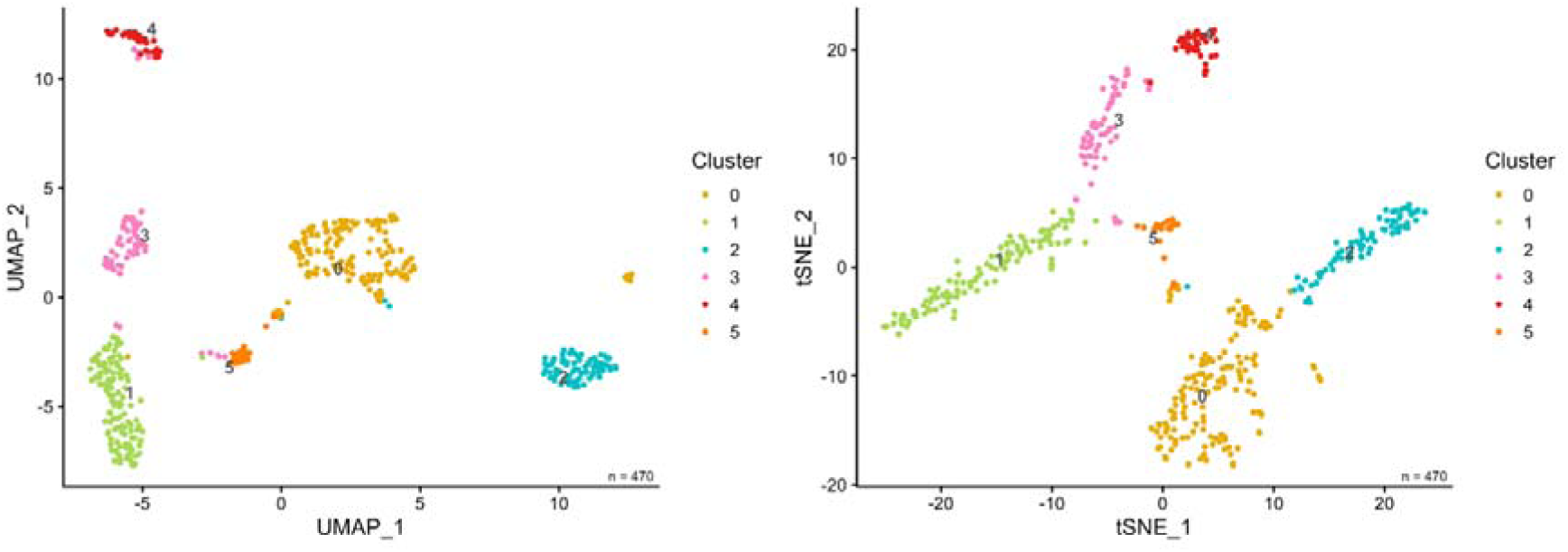
t-SNE embeddings of the filtered single-cell dataset. t-SNE projections of the filtered neural complex dataset with cells colored by cluster and labeled with the final biological cluster names.

**Figure S3.**
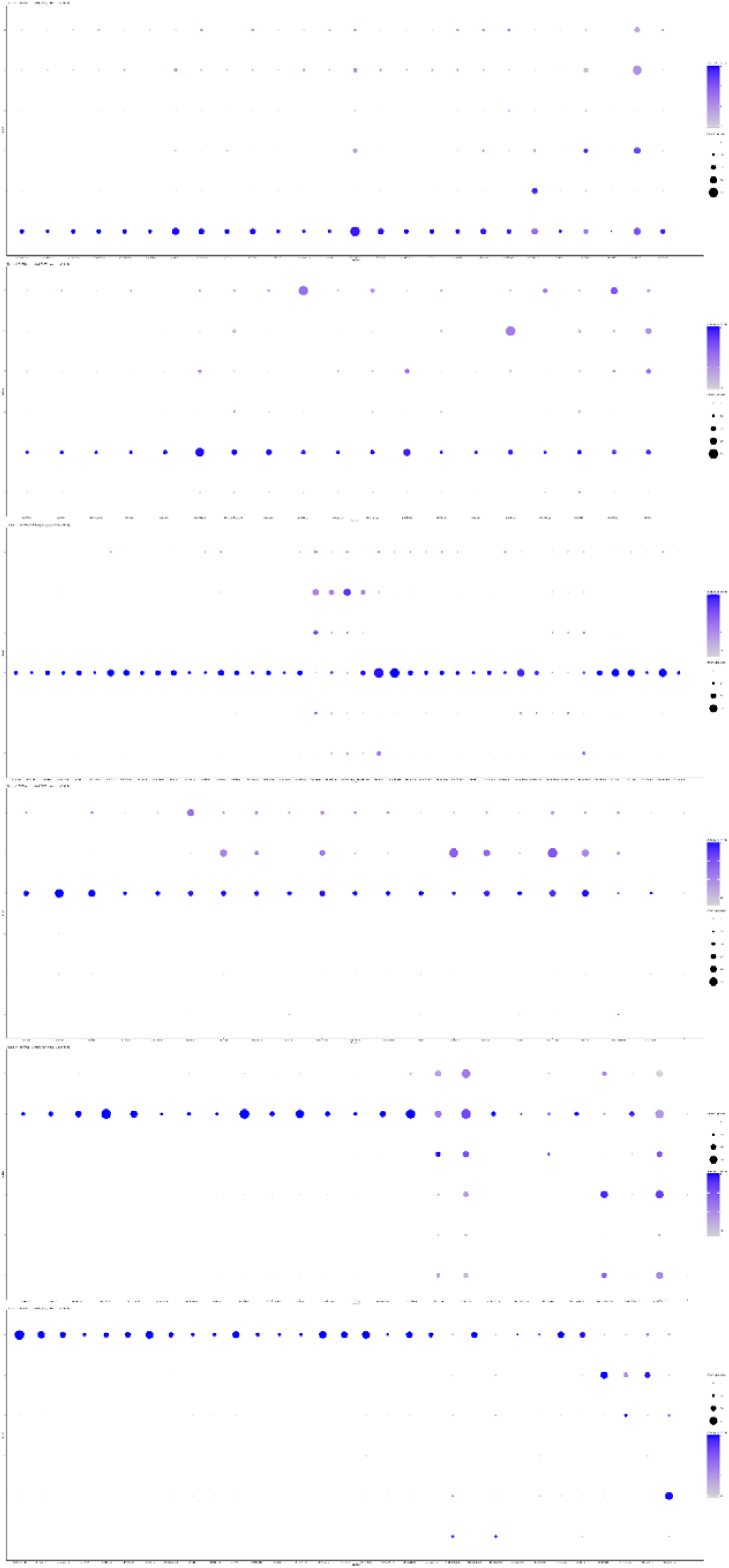
Extended marker visualization across clusters. Dot plot showing filtered marker sets for each cluster, including top-ranked differentially expressed genes and curated anchor markers used during cluster annotation. Genes expressed in less than 10% of cells in the target cluster were excluded from the visualization for display purposes only. Dot size represents the fraction of cells expressing each gene within a cluster, and color intensity reflects average expression.

**Figure S4A.**
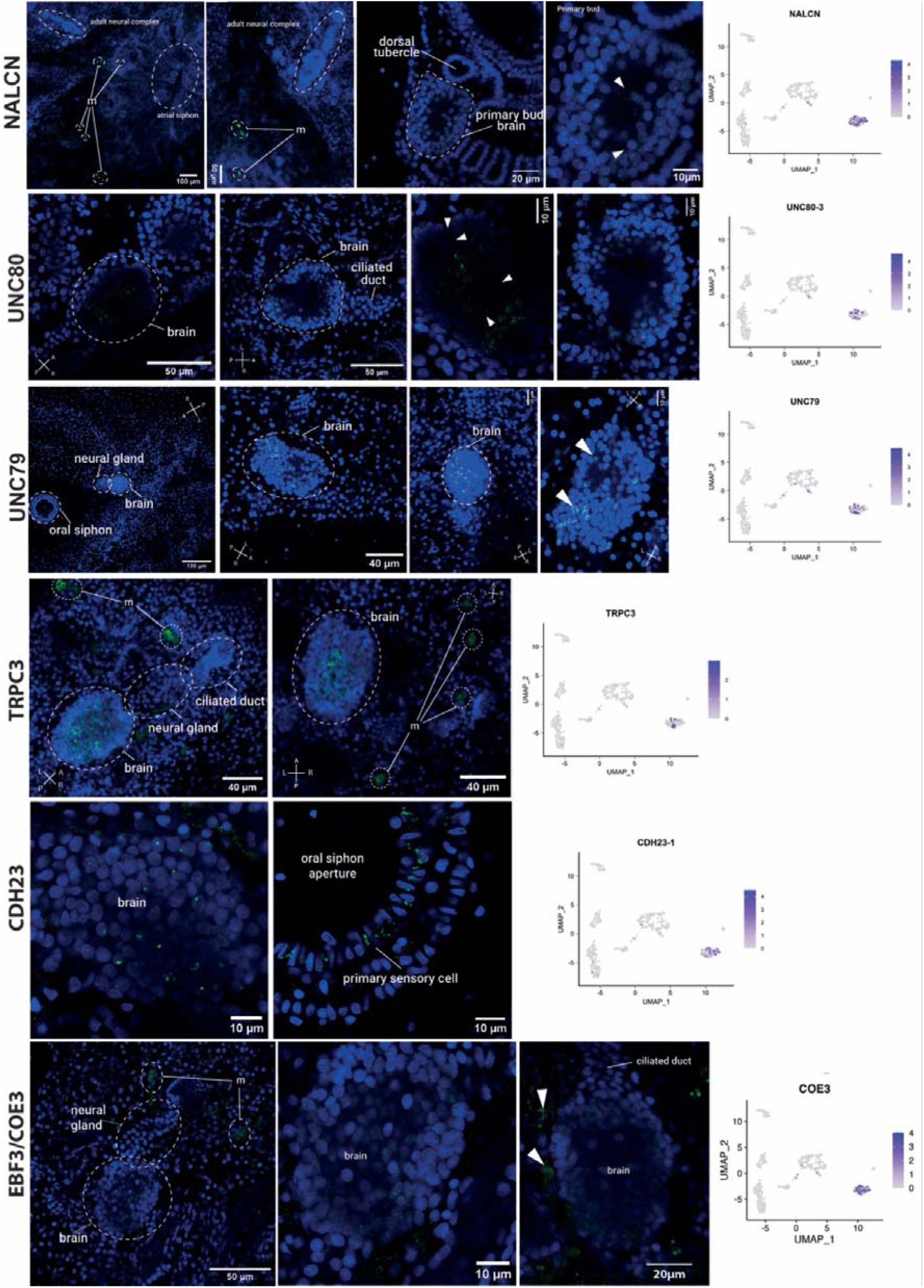
Extended HCR-FISH validation of cluster 2-associated genes. Representative HCR-FISH images and corresponding UMAP feature plots for selected genes associated with the neuronal cluster of the Smart-seq3 adult neural-complex atlas, including NALCN, UNC80-3, UNC79, TRPC3, CDH23-1, and EBF3/COE3. For each gene, the UMAP feature plot shows normalized expression across all cells in the single-cell dataset, with color intensity indicating log-normalized expression levels, whereas the HCR-FISH images show the spatial distribution of the corresponding transcripts in adult zooids and developing buds. Green indicates HCR-FISH signal and blue indicates Hoechst/DAPI nuclear counterstain. Dashed outlines indicate anatomical structures, including the brain, neural gland, ciliated duct, dorsal tubercle, primary bud brain, and oral siphon when visible. Arrowheads indicate HCR-positive cells or expression domains. Scale bars are indicated in each panel. NALCN, UNC80-3, and UNC79 show enrichment in the neuronal cluster in the single-cell atlas and localize to sparse cells associated with the adult brain, neural-complex-adjacent territories, and developing bud neural structures. NALCNsignal is also detected in the developing primary bud brain. UNC80-3 and UNC79 are detected in cells within or adjacent to the adult brain and nearby neural-complex-associated domains, consistent with a neuronal/excitability-related identity. TRPC3 is enriched in the neuronal cluster and is detected in cells within the adult brain as well as in adjacent neural-complex-associated territories, including the neural gland/ciliated duct region, with additional scattered mantle-associated cells (m) surrounding the neural complex. CDH23-1 is enriched in the neuronal cluster and is detected both in the adult brain region and in peripheral sensory-associated cells around the oral siphon aperture, including primary sensory cells. EBF3/COE3 is enriched in the neuronal cluster and is detected in the adult brain and in additional cells associated with the neural gland/ciliated duct region and surrounding territories, consistent with a broader neuronal differentiation-associated expression domain.

**Figure S4B.**
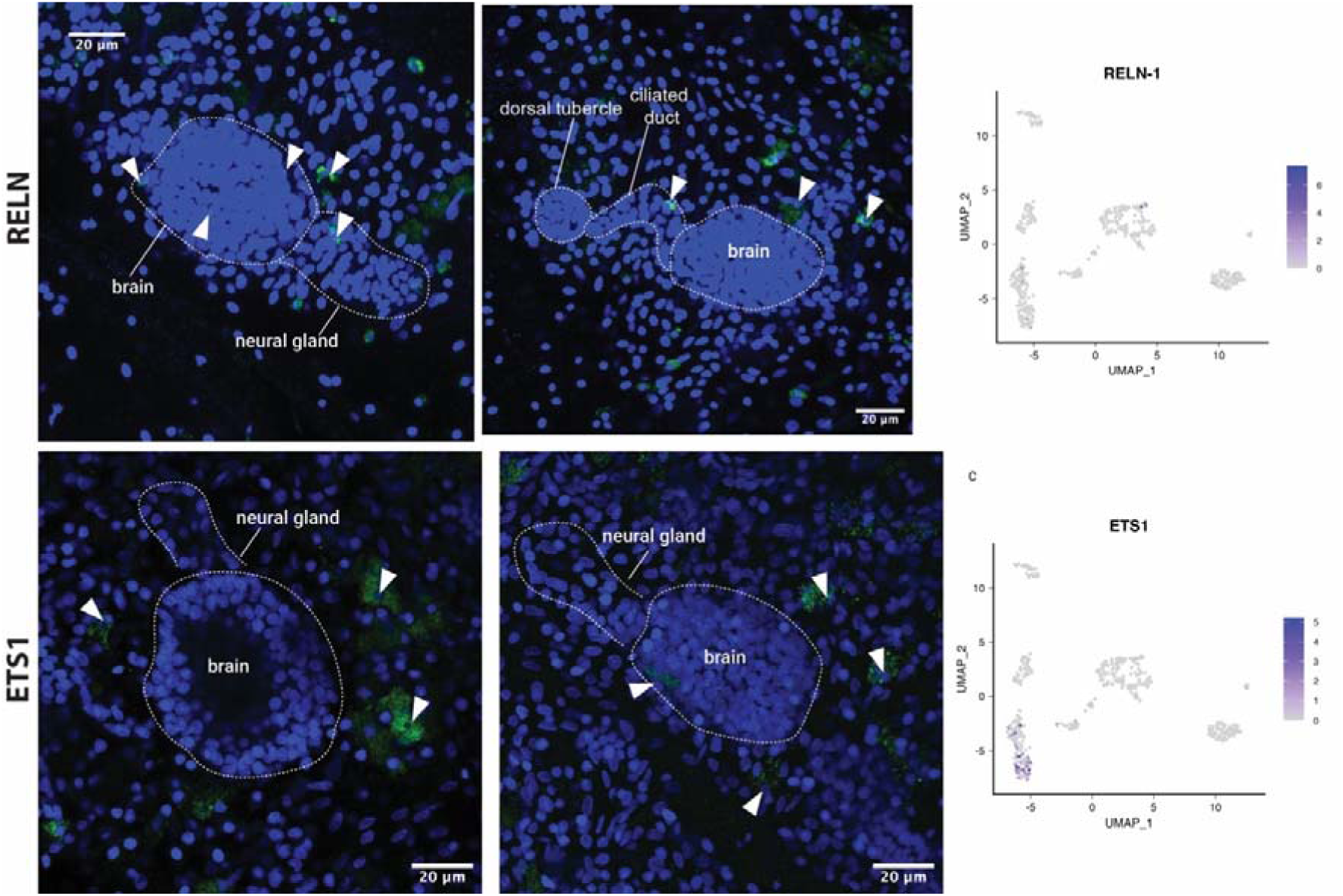
Extended HCR-FISH validation of cluster 1-associated genes. Representative HCR-FISH images and corresponding UMAP feature plots for RELN-1 and ETS1, two genes associated with cluster 1 in the Smart-seq3 single-cell atlas. For each gene, the UMAP feature plot shows normalized expression across all cells of the adult *Botryllus schlosseri* neural-complex dataset, with color intensity indicating log-normalized expression levels. HCR-FISH images show the spatial distribution of the corresponding transcripts in adult zooids and neural-complex-associated territories. RELN-1 signal is detected in sparse cells associated with the adult brain/neural gland region, the dorsal tubercle/ciliated duct area, and surrounding brain-adjacent tissues. ETS1 signal is detected in scattered cells surrounding the adult brain and neural gland region, as well as in additional cells distributed in adjacent adult zooid tissues. Green indicates HCR-FISH signal and blue indicates Hoechst/DAPI nuclear counterstain. Dashed outlines indicate anatomical structures, including the brain, neural gland, dorsal tubercle, and ciliated duct when visible. Arrowheads indicate HCR-positive cells or expression domains. Scale bars are indicated in each panel.

**Figure S4C.**
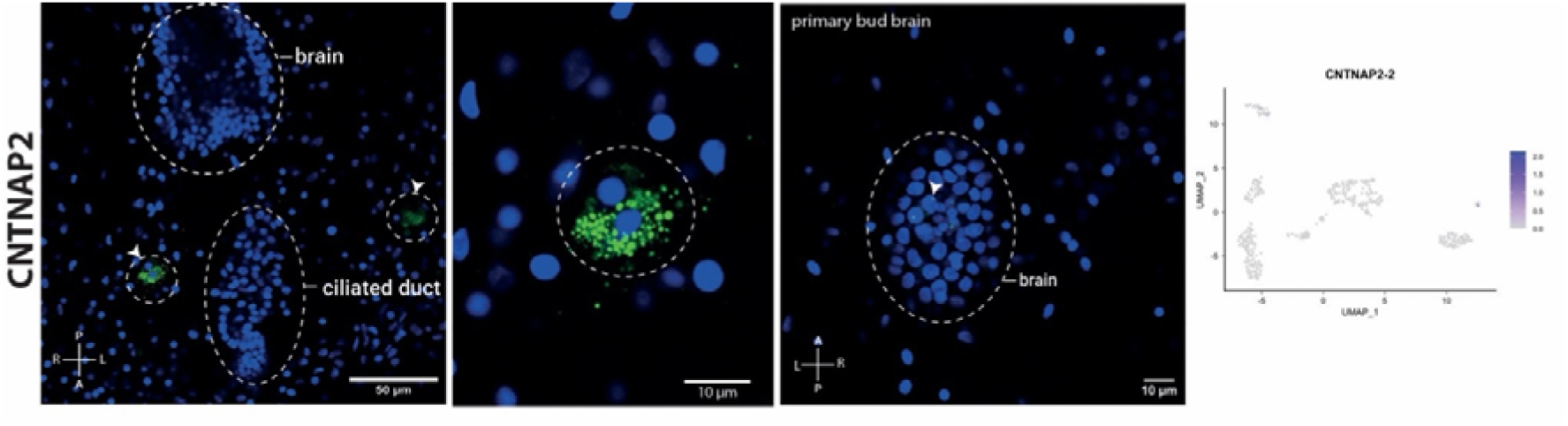
HCR-FISH validation of CNTNAP2-2, cluster 4. Representative HCR-FISH images and corresponding UMAP feature plot for CNTNAP2-2. In the adult zooid, CNTNAP2-2 signal is detected in discrete cell clusters located in mantle/perineural territories adjacent to the brain and ciliated duct. Higher magnification shows cytoplasmic CNTNAP2-2 signal in closely associated cells. In the primary bud, CNTNAP2-2-positive cells are detected in the developing bud brain. The paired UMAP feature plot shows restricted expression of CNTNAP2-2 in a small subset of cells in the Smart-seq3 atlas. Green, HCR-FISH signal; blue, DAPI/Hoechst nuclear counterstain. Dashed outlines indicate anatomical structures, and arrowheads indicate positive cells or expression domains. Scale bars are indicated in each panel.

**Figure S4D.**
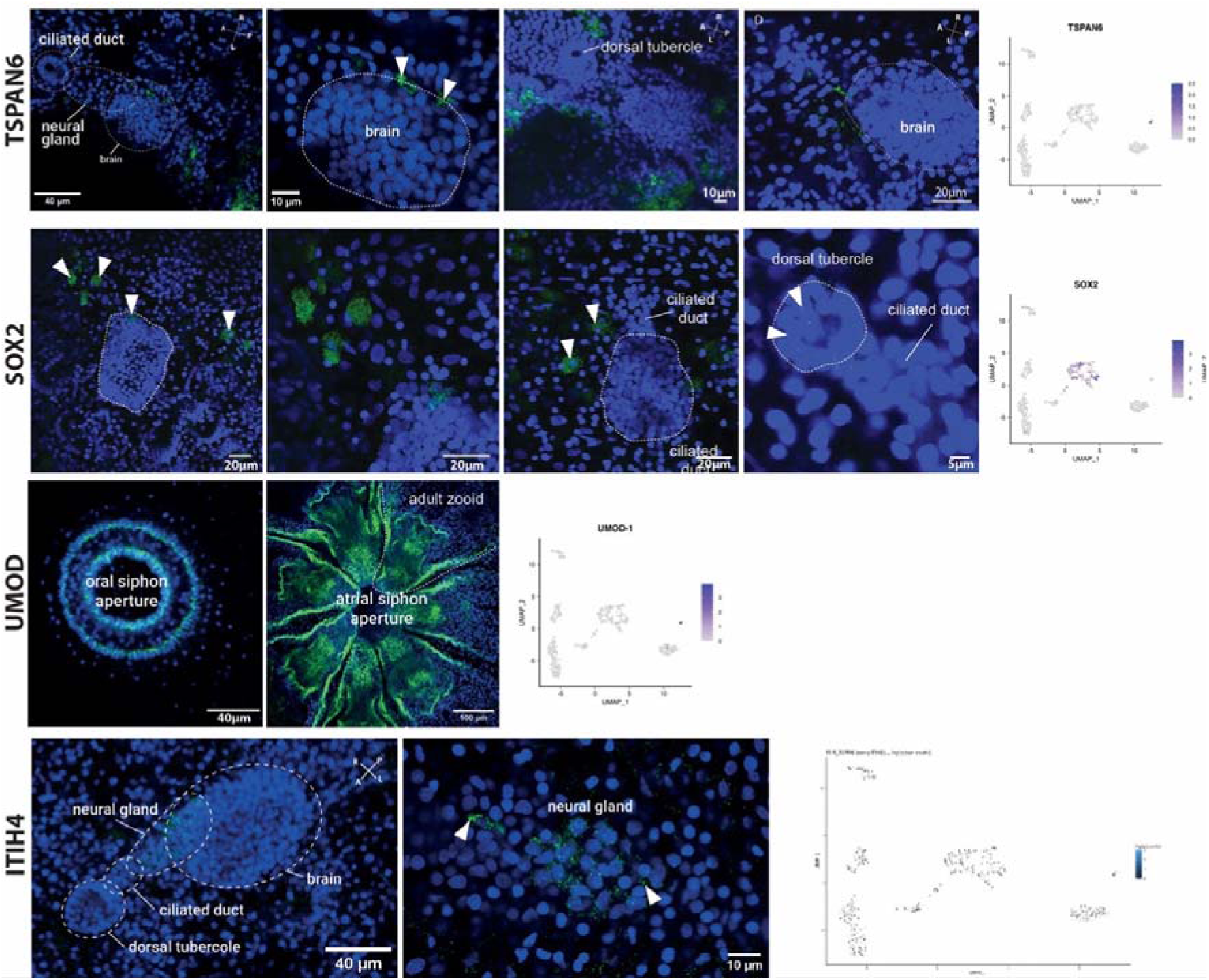
Extended HCR-FISH validation of cluster 0-associated genes. Representative HCR-FISH images and corresponding UMAP feature plots for selected genes associated with the neuroepithelial/progenitor-like transcriptional programs, including TSPAN6, SOX2, UMOD-1, and ITIH4. For each gene, UMAP feature plots show normalized expression across the Smart-seq3 single-cell atlas, with color intensity indicating log-normalized expression levels. HCR-FISH images show the spatial distribution of the corresponding transcript in adult zooids and neural-complex-associated tissues. Green indicates HCR-FISH signal and blue indicates Hoechst or DAPI nuclear counterstain. Dashed outlines indicate anatomical structures, including the brain, neural gland, ciliated duct, and dorsal tubercle when visible. Arrowheads indicate HCR-positive cells or expression domains. Scale bars are indicated in each panel. TSPAN6 expression is detected in cells surrounding the adult brain and in adjacent perineural or mantle-associated territories, consistent with its expression in the single-cell atlas. SOX2, which is enriched in the neuroepithelial/progenitor-like cluster, is detected in cells surrounding the adult brain, in adjacent mantle/perineural territories, and weakly in the ciliated duct region. UMOD-1 shows limited expression in the single-cell atlas and spatially labels extra-brain epithelial territories, particularly siphonal domains including the oral and atrial siphon apertures. ITIH4 expression is detected in the neural gland region; higher magnification shows cytoplasmic ITIH4 signal in individual neural-gland-associated cells

**Figure S4E:**
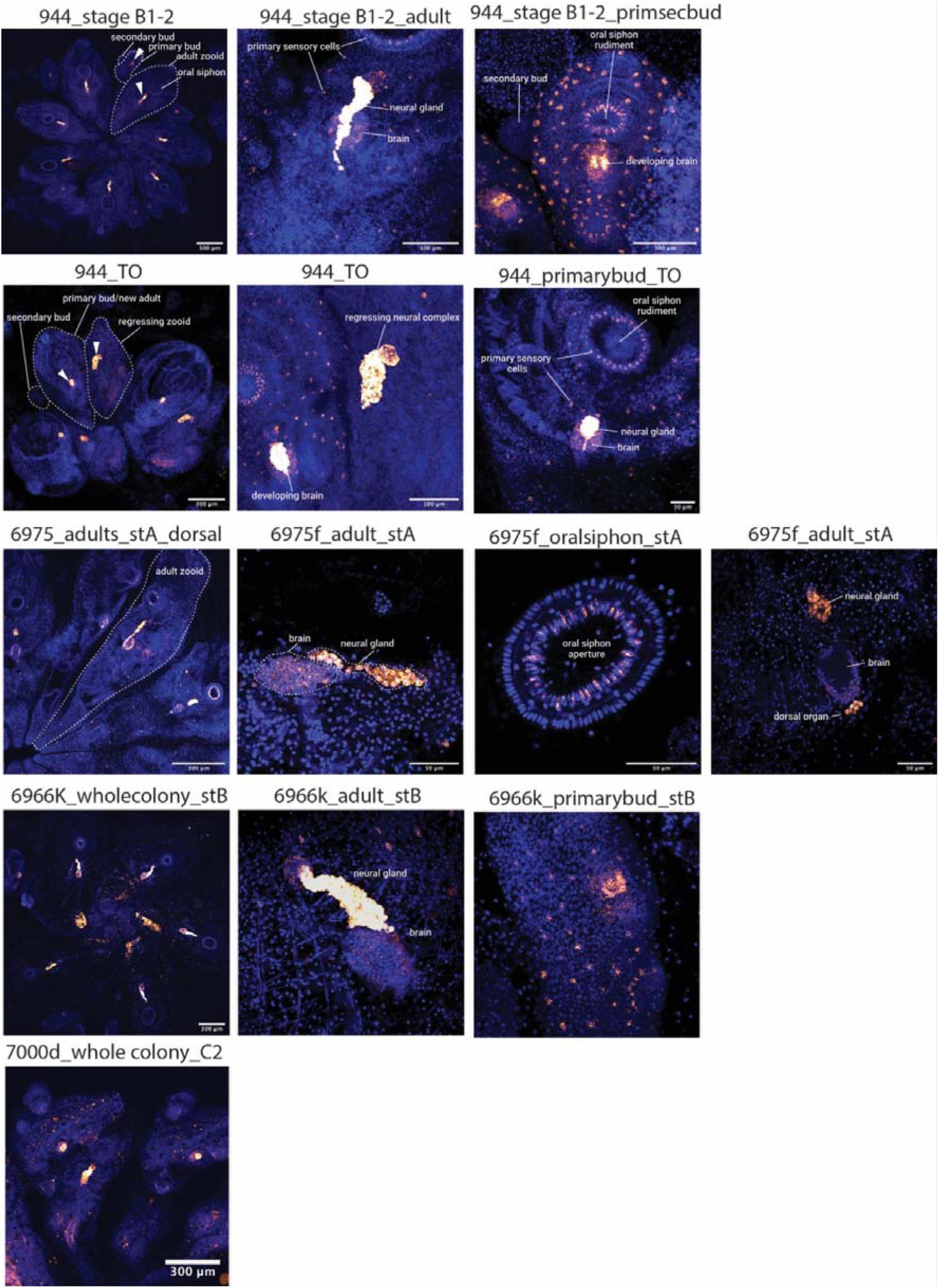
Spatial expression of APP across blastogenetic stages and neural-complex-associated territories. HCR-FISH detection of APP transcripts reveals a broad and dynamic expression pattern across adult zooids and developing buds. In stage B colonies, APP signal is detected in the neural complex of the adult zooid as well as in the developing brain of the primary bud. During takeover (TO), APP expression is observed in both the regressing adult neural complex and in the developing neural complex of the primary bud within the same colony, highlighting its presence across degenerative and developmental phases.In stage A adults, APP expression is particularly prominent in the neural gland and ciliated duct, where signal intensity exceeds that observed in the cerebral ganglion, which remains positively labeled but at lower levels. Across all stages, APP is consistently detected in primary sensory cells, corresponding to mechanoreceptive neurons distributed in the mantle epithelium and surrounding the oral siphon. At later stages (C2), APP expression is maintained in the neural complex of adult zooids as well as in primary and secondary buds.

**Figure S4F:**
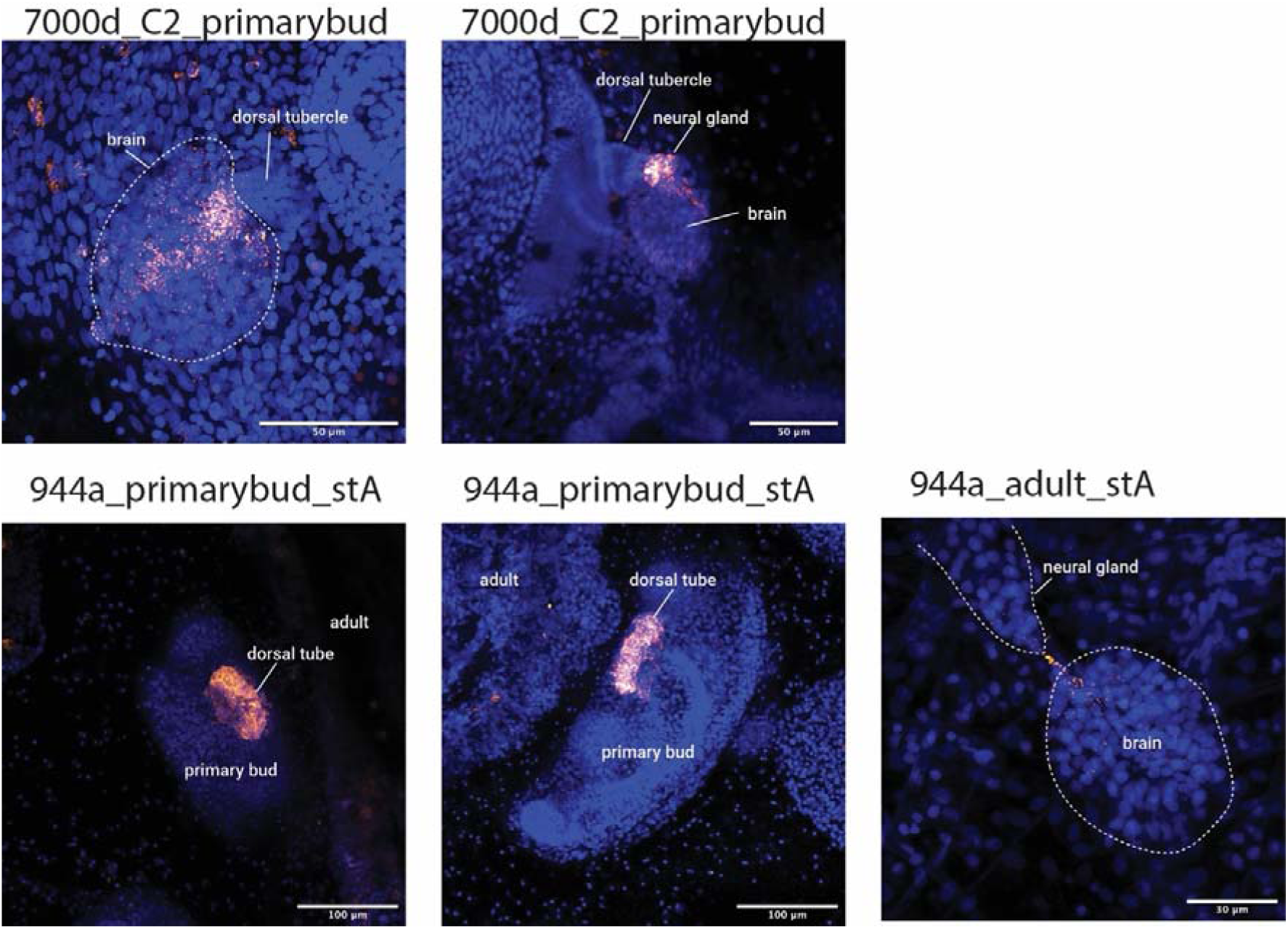
Spatial expression of NEUROG1 in developing and adult neural-complex-associated territories in *Botryllus schlosseri*. HCR-FISH detection of NEUROG1 reveals a stage-dependent and spatially restricted expression pattern associated with neural-complex development. In primary buds at stage C2, NEUROG1 expression is detected within the neural complex, with stronger signal in the neural gland region compared to the developing cerebral ganglion. At earlier stages (stage A), NEUROG1 prominently marks the dorsal tube of the primary bud, a transient neurogenic structure that gives rise to both the neural gland and the cerebral ganglion. In adult zooids (stage A), NEUROG1 expression is markedly reduced, with only weak signal detected in the cerebral ganglion and a small number of positive cells located between the brain and the neural gland.

**Figure S4G.**
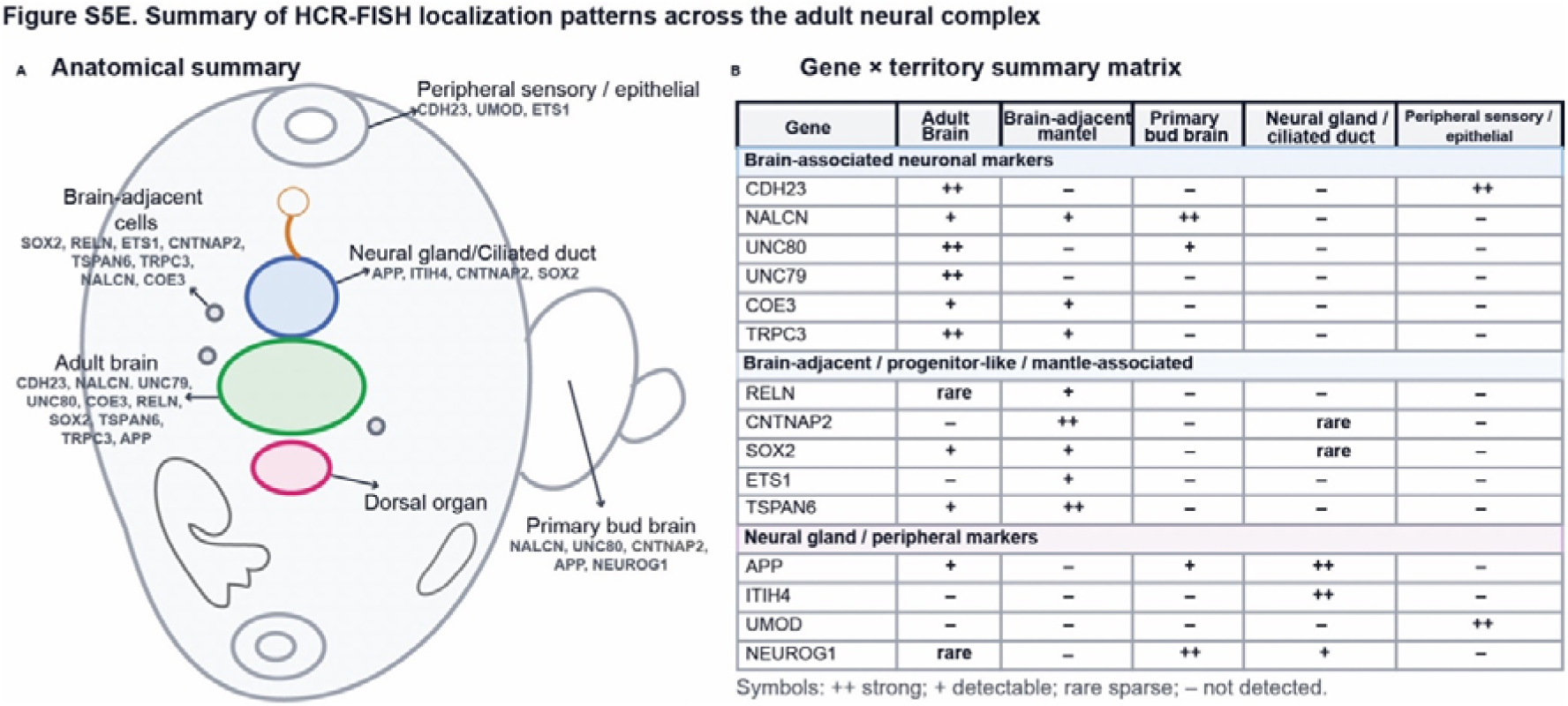
Summary of HCR-FISH localization patterns across the adult *B. schlosseri* neural complex. (A) Schematic overview of the principal anatomical territories labeled by selected HCR-FISH markers in the adult zooid and neural-complex-associated region. Marker expression resolves multiple spatial categories, including adult brain-associated neuronal domains, brain-adjacent mantle-associated territories, neural gland/ciliated duct-associated domains, primary bud neural territories, and peripheral sensory/epithelial regions. (B) Matrix summarizing the predominant localization pattern of each HCR marker across major anatomical domains. Symbols indicate strong (++), detectable (+), sparse (rare), or undetected (**–**) signal in each territory.

**Figure S5.**
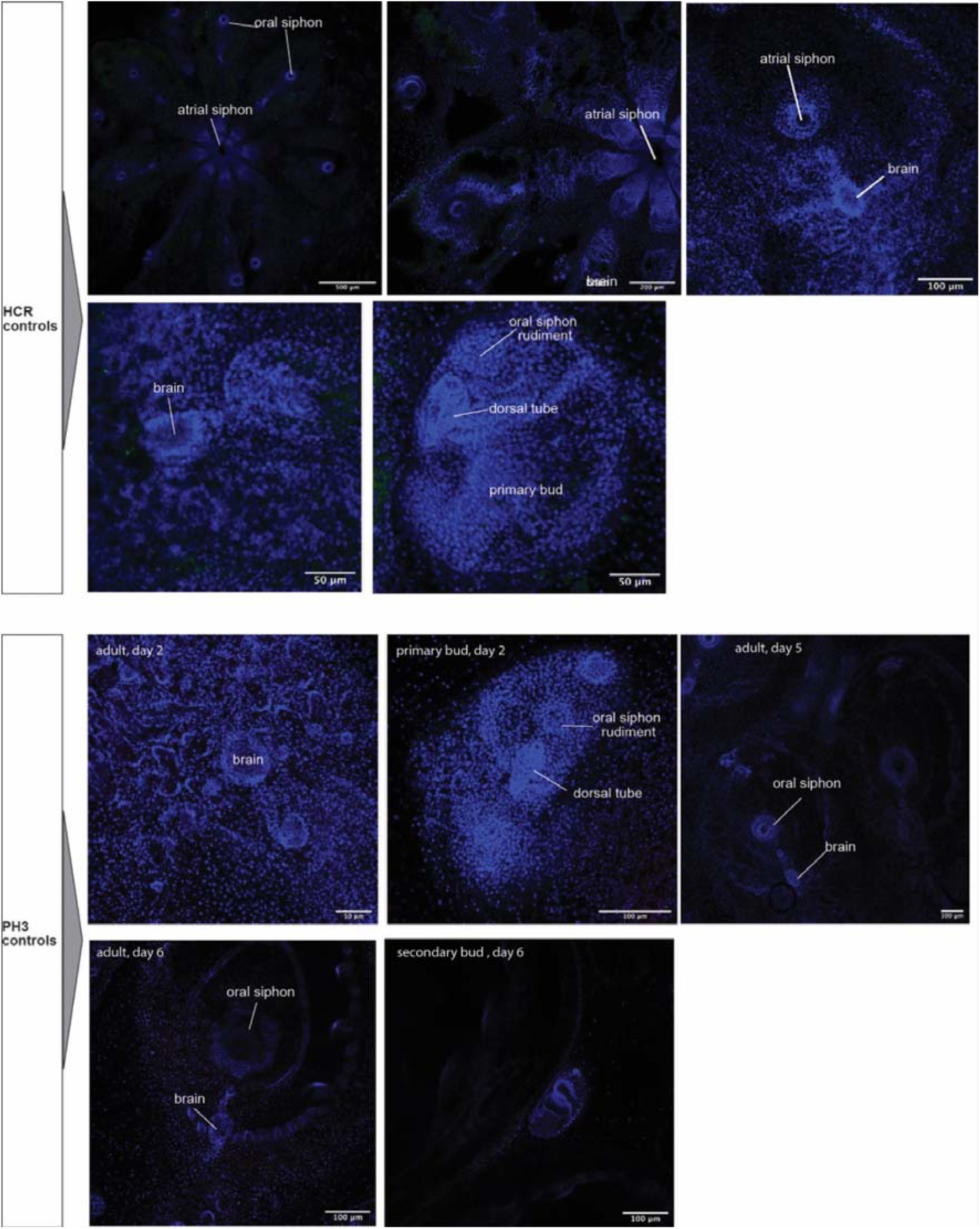
Negative controls for HCR-FISH and PH3 immunostaining. Representative control images showing background signal in the absence of specific HCR probe signal or PH3 immunoreactivity. The upper panels show HCR-FISH control samples across adult zooids and developing buds, including views of the adult brain, atrial and oral siphons, and the primary bud dorsal tube. The lower panels show PH3 immunostaining controls in adult zooids and buds at different days of the blastogenetic cycle, including adult Day 2, primary bud Day 2, adult Day 5, adult Day 6, and secondary bud Day 6. Nuclei are counterstained with DAPI in blue. Anatomical landmarks are indicated to show the position of the brain/neural complex, oral and atrial siphons, oral siphon rudiment, dorsal tube, and buds.. Scale bars are indicated in each image.

**Figure S6.**
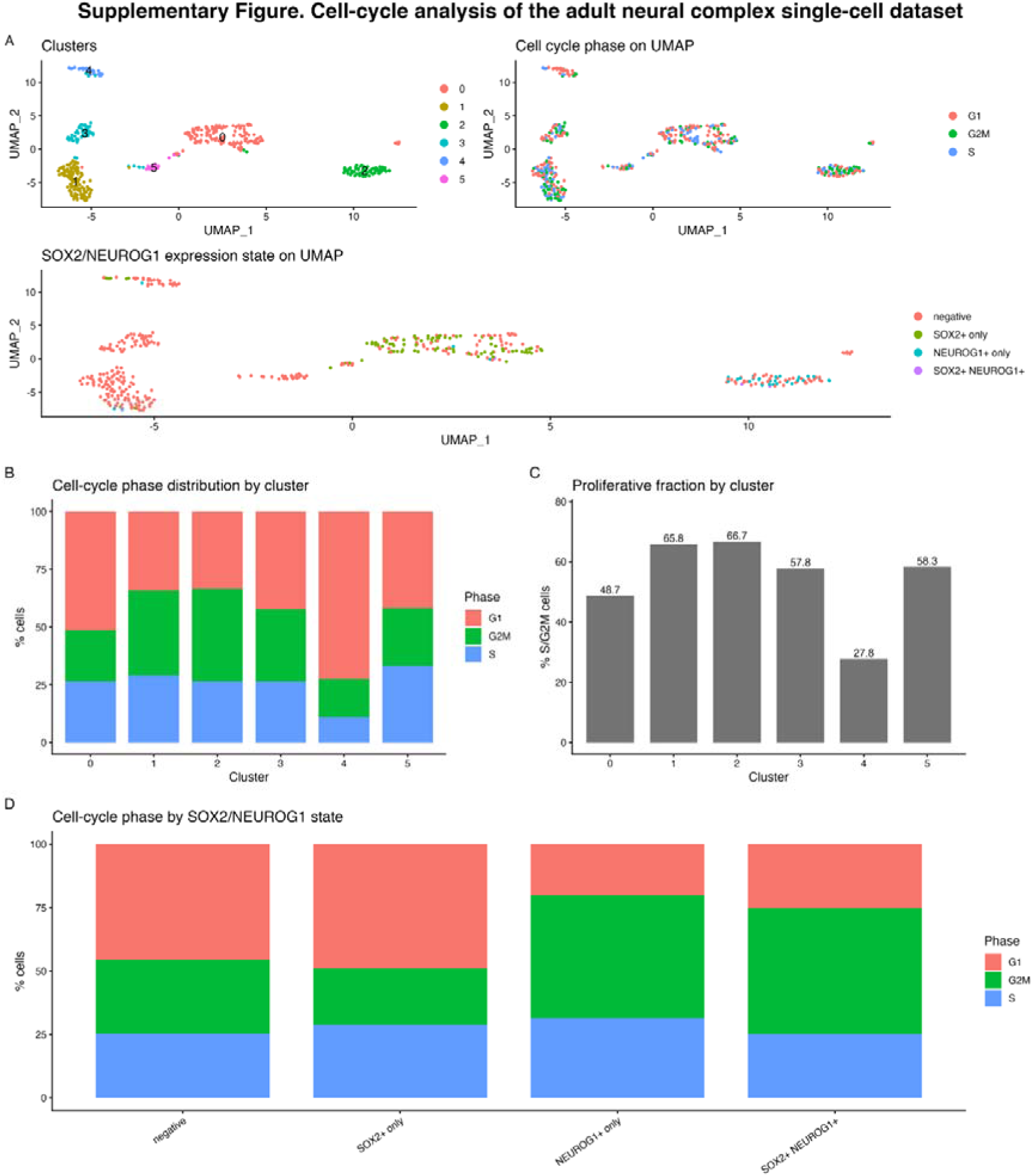
Cell-cycle phase distribution across clusters. (A) UMAP representation of the single-cell dataset showing cluster identity, cell-cycle phase assignment, and SOX2/NEUROG1 expression state. SOX2-positive cells are mainly localized in cluster 0, whereas NEUROG1-positive cells are concentrated in cluster 2. (B) Distribution of cells assigned to G1, S, or G2/M phases across clusters. (C): Proliferative fraction by cluster, calculated as the percentage of cells assigned to S or G2/M phases. Clusters 1 and 2 showed the highest S/G2/M fractions, whereas cluster 4 showed the lowest proliferative fraction. (D): Cell-cycle phase distribution stratified by SOX2/NEUROG1 expression state. SOX2-only cells showed a mixed G1 and S/G2/M profile, while NEUROG1-only cells were predominantly assigned to S/G2/M phases. SOX2/NEUROG1 double-positive cells were rare and should be interpreted cautiously. Overall, these analyses support the presence of actively cycling progenitor-like and neurogenic/committed precursor-like populations within the adult neural complex.

**Figure S6B.**
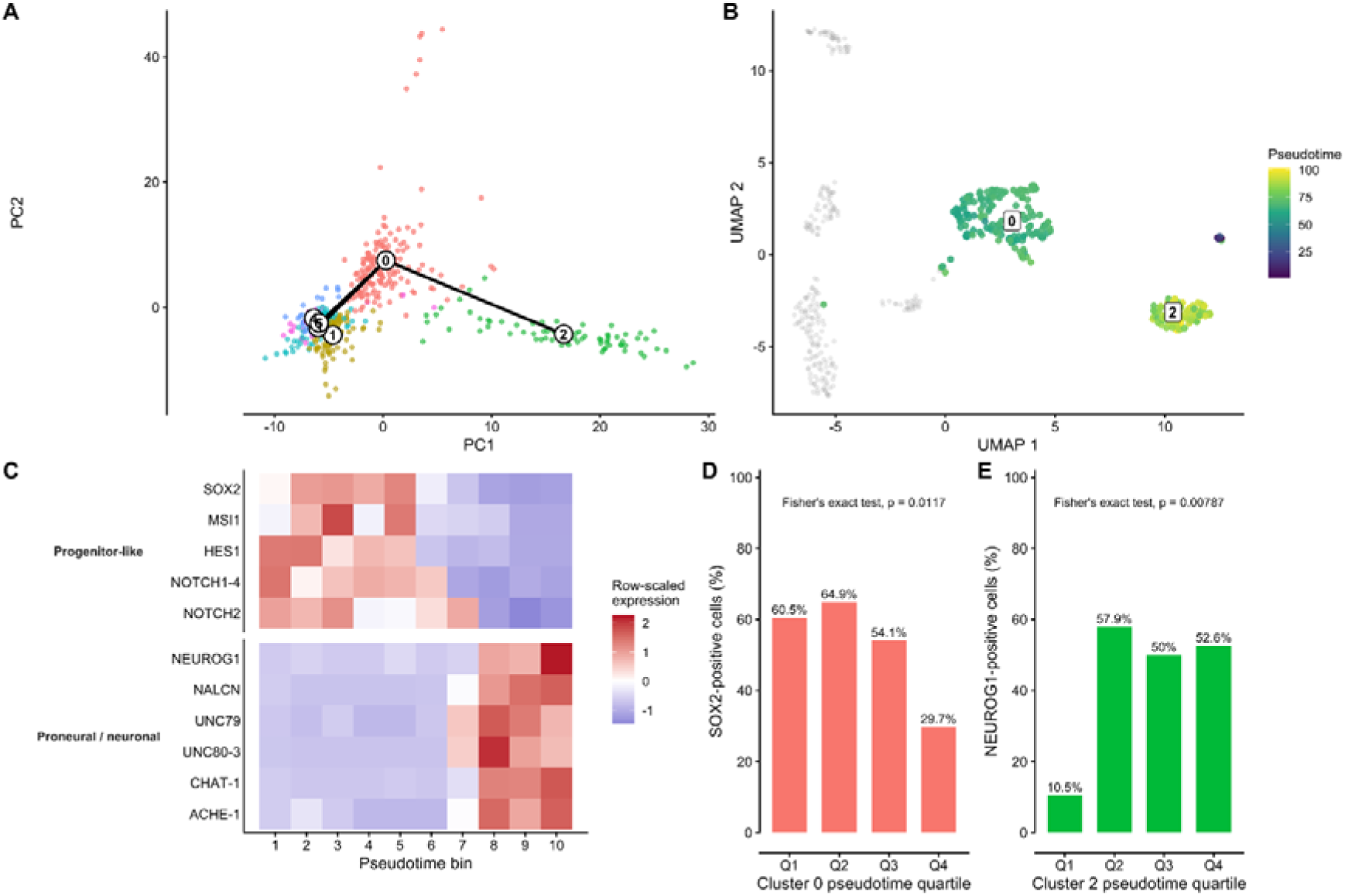
Trajectory inference identifies a transcriptional axis linking progenitor-like and neuronal states. (A) Unrooted Slingshot trajectory inference based on the first five principal components of the single-cell dataset. The cluster-level topology is displayed in the PC1–PC2 projection and directly connects progenitor-like cluster 0 with neuronal cluster 2 before specification of a starting state. (B) Pseudotime along the inferred lineage connecting clusters 0 and 2 after cluster 0 was specified as a biologically informed starting state. Pseudotime was inferred in five-dimensional PCA space and projected onto the existing UMAP for visualization. The lineage contained 149 of 154 cluster 0 cells and all 75 cluster 2 cells; cells not assigned to this lineage are shown in gray. (C) Expression dynamics of selected progenitor-associated and proneural/neuronal genes along the inferred 0–2 transcriptional axis. Cells were divided into ten equal-frequency pseudotime bins, mean normalized expression was calculated within each bin, and expression values were scaled independently for each gene across bins. (D) Fraction of *SOX2*-positive cells within cluster 0 across pseudotime quartiles. SOX2-positive cells represented 60.5%, 64.9%, 54.1%, and 29.7% of cells in Q1– Q4, respectively (n = 38, 37, 37, and 37). SOX2 positivity differed significantly across quartiles (Fisher’s exact test, p = 0.0117). Pairwise proportion tests with Benjamini– Hochberg correction showed significant reductions in Q4 relative to Q1 (adjusted p = 0.043) and Q2 (adjusted p = 0.031). (E) Fraction of *NEUROG1*-positive cells within cluster 2 across pseudotime quartiles. *NEUROG1*-positive cells represented 10.5%, 57.9%, 50.0%, and 52.6% of cells in Q1–Q4, respectively (n = 19, 19, 18, and 19). *NEUROG1* positivity differed significantly across quartiles (Fisher’s exact test, p = 0.00787). Pairwise proportion tests with Benjamini–Hochberg correction showed significantly lower *NEUROG1* positivity in Q1 than in Q2 (adjusted p = 0.037), Q3 (adjusted p = 0.047), and Q4 (adjusted p = 0.044).

**Figure S7:**
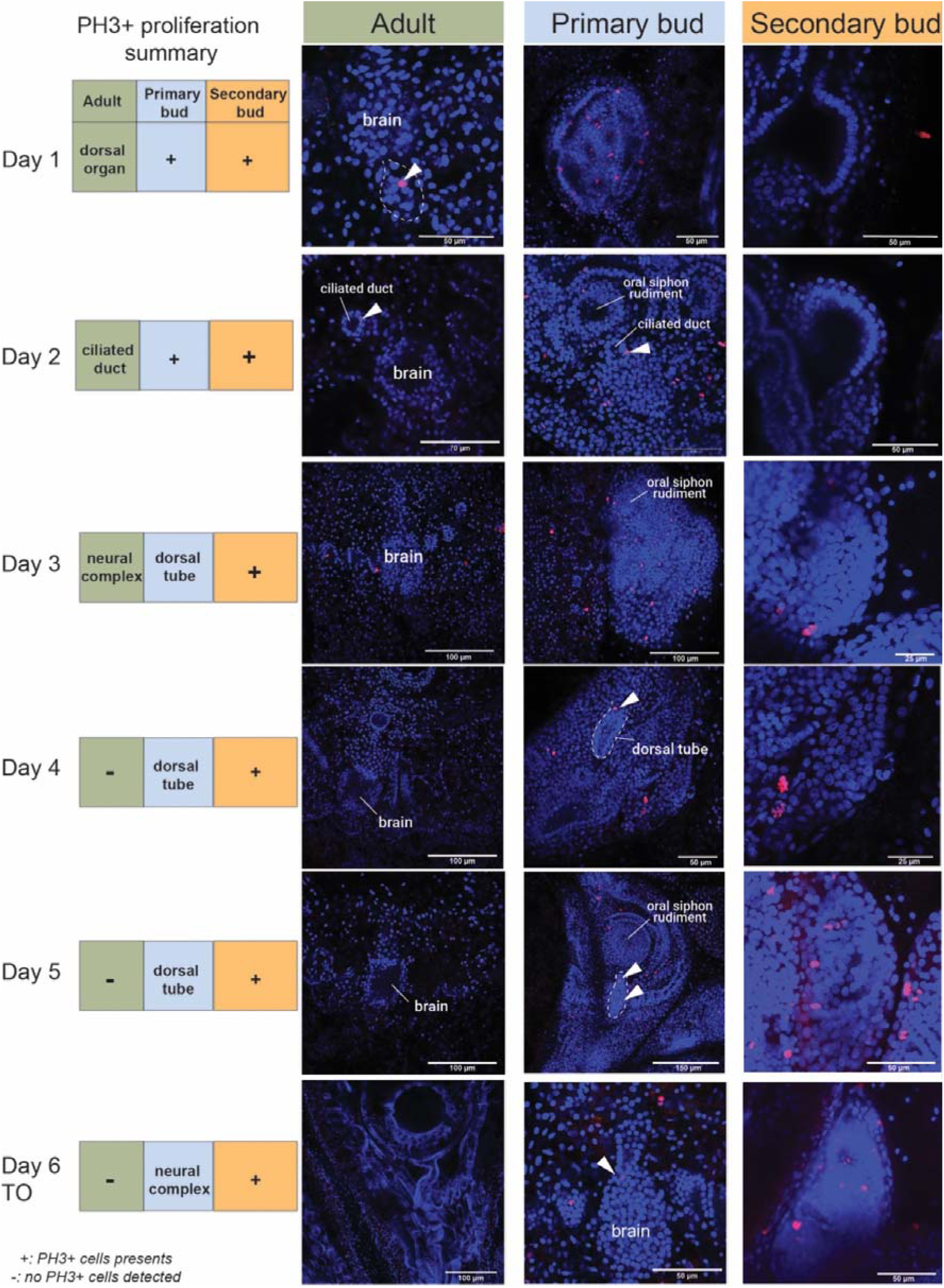
Stage-dependent PH3 labeling in the neural complex of adult and developing zooids across the blastogenetic cycle. Representative confocal images of phospho-Histone H3 (PH3) immunostaining across successive blastogenetic stages, showing the adult, primary bud, and secondary bud neural regions from Day 1 to Day 6/takeover (TO). PH3-positive cells are detected in red and nuclei are counterstained in blue. The summary table on the left indicates, for each stage and generation, the anatomical region in which PH3+ cells were detected when this could be assigned with confidence. In the adult, PH3+ cells are observed mainly at early stages and are associated with the dorsal organ or ciliated duct region, whereas they are absent at intermediate late stages. In buds, proliferative cells are detected throughout development, although in early stages the signal cannot always be assigned unambiguously to a defined neural substructure. “+” indicates the presence of PH3+ cells without clear anatomical assignment; “−” indicates no PH3+ cells detected. Together, these data support persistent proliferative activity in developing buds and a more spatially restricted, stage-dependent pattern in the adult neural complex.

**Figure S8.**
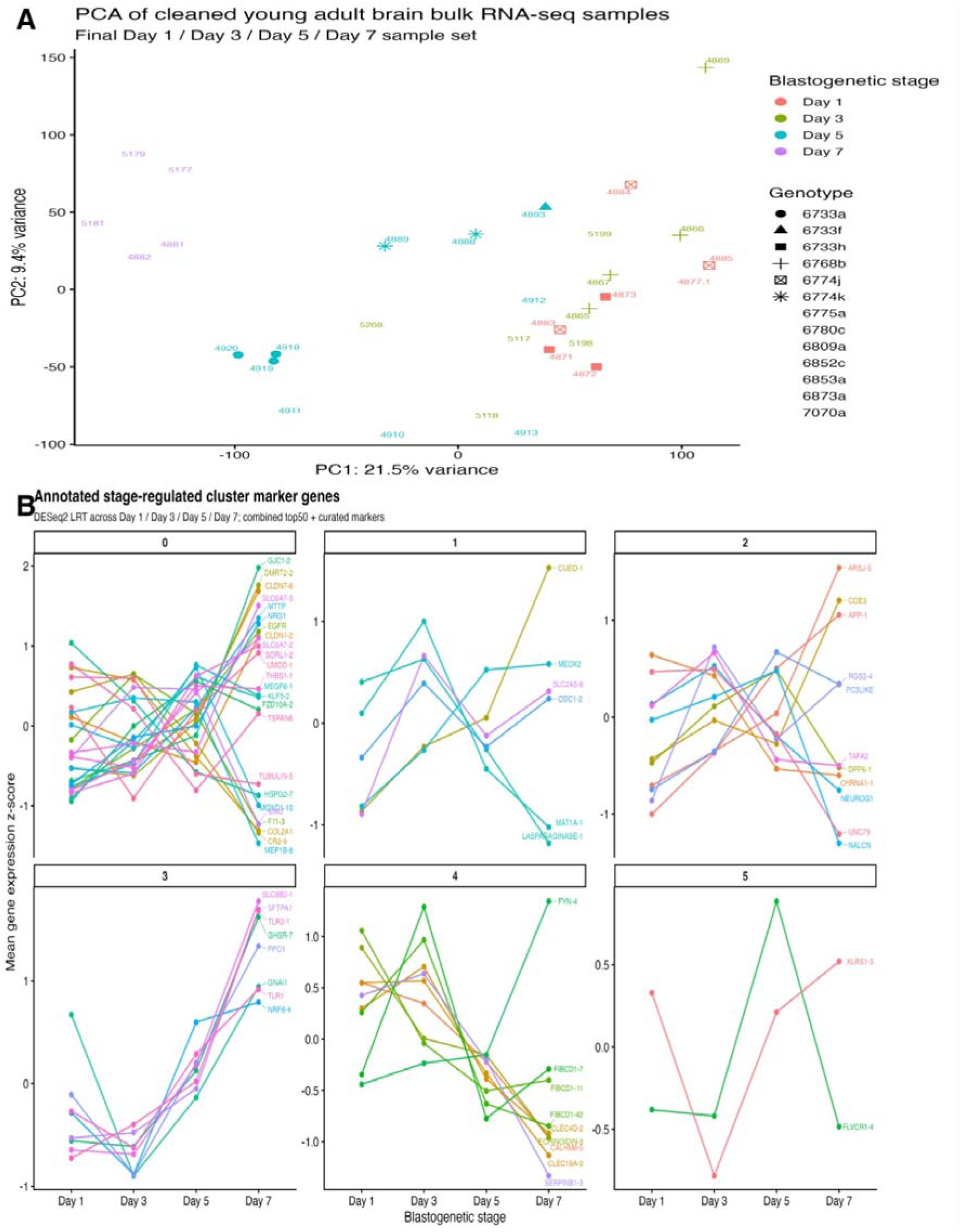
Additional bulk RNA-seq analyses across the blastogenetic cycle. (A) Principal component analysis of the cleaned young adult *Botryllus schlosseri* brain bulk RNA-seq dataset. Samples correspond to Day 1, Day 3, Day 5, and Day 7 of the blastogenetic cycle, corresponding to stages A, B, C, and D, respectively. Each point represents one bulk brain sample. Colors indicate blastogenetic stage and symbols indicate genotype. The PCA shows the global transcriptional structure of the cleaned dataset after removal of low-quality or misclassified samples. Later-stage samples, particularly Day 5 and Day 7, show partial separation from early Day 1/Day 3 samples along the first two principal components, consistent with stage-associated transcriptional variation. (B) Annotated stage-regulated marker genes grouped by single-cell-derived cluster. Genes were selected by intersecting bulk stage-regulated genes identified by DESeq2 likelihood-ratio testing across Day 1, Day 3, Day 5, and Day 7 with the combined single-cell marker list. Only annotated genes were retained for visualization. Each line represents one gene, shown as mean z-scored expression across stages. Gene labels are shown at the final stage to facilitate identification of the major stage-associated trends. This panel provides an expanded view of the cluster-associated genes that vary significantly across the blastogenetic cycle.

**Figure S9.**
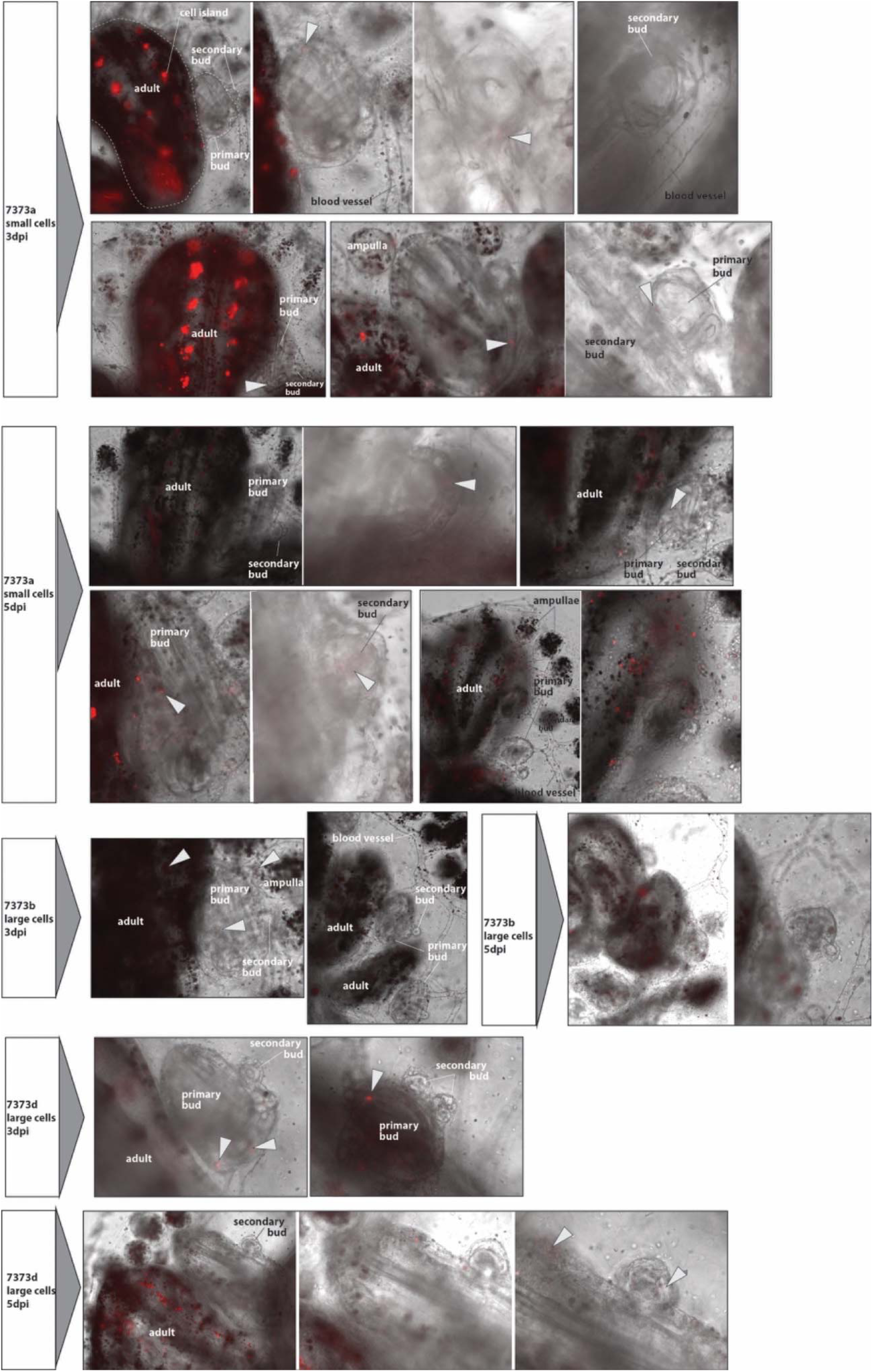
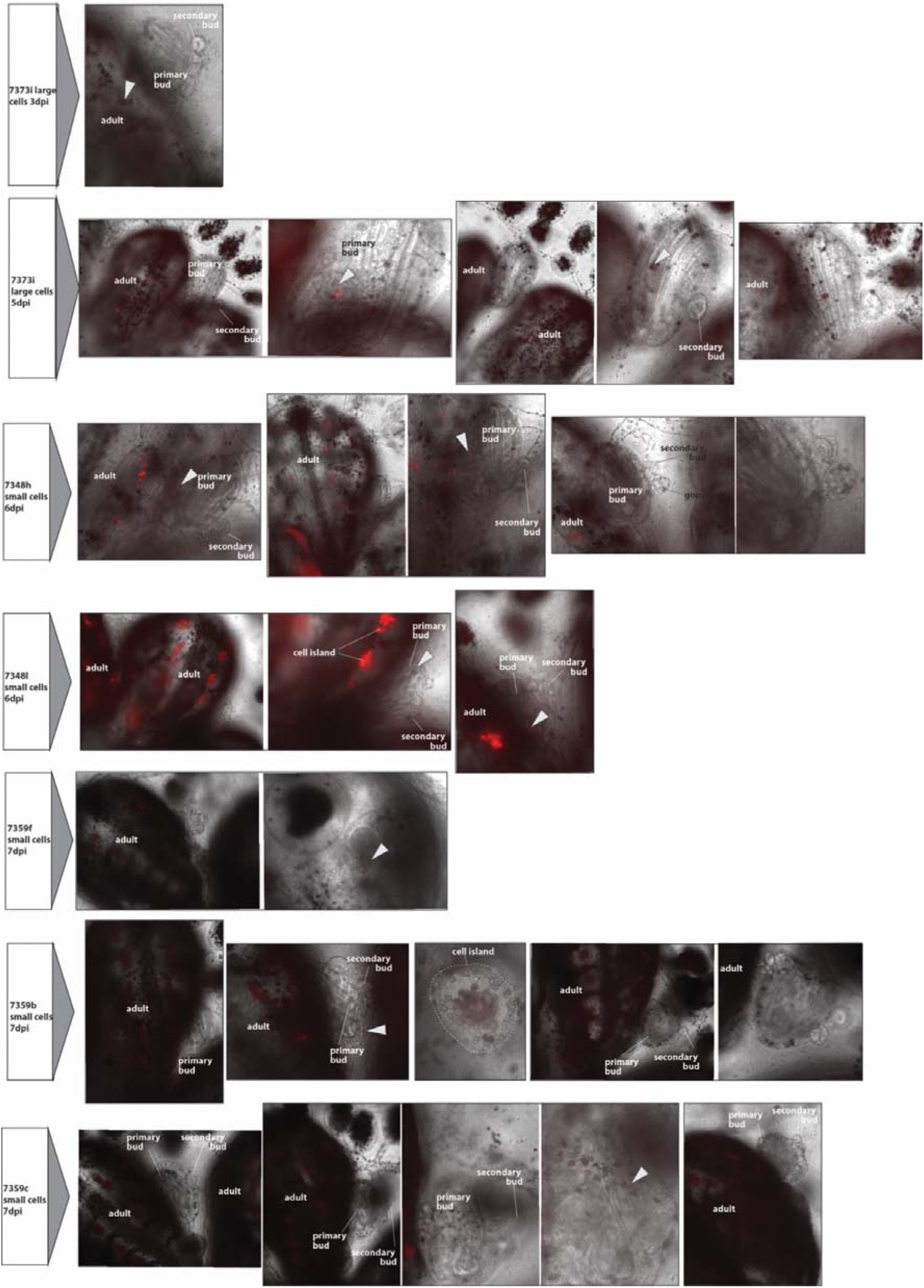
*In vivo* tracking of DiD-labelled cells following transplantation into recipient *Botryllus schlosseri*colonies. Representative live-imaging fields documenting the distribution of fluorescently labelled cells after injection of FACS-isolated small-or large-cell fractions into the extracorporeal vasculature of recipient colonies. DiD fluorescence is shown in red and is overlaid on transmitted-light images to visualize the position of labelled cells relative to the adult zooid, primary and secondary buds, peripheral ampullae, blood vessels, endostyle-associated regions, gonadal territories, and circulating or tissue-associated cell islands. Images are grouped according to the injected cell fraction and post-injection observation time, as defined in the experimental dataset. White arrowheads indicate representative DiD-positive cells or cell aggregates, while dashed outlines identify selected anatomical structures or cell islands where shown. Labelled cells were detected in adult zooids and developing buds, as well as in the shared vascular system and peripheral ampullae, with substantial variability among colonies, anatomical compartments, and imaging fields. The full image series is provided to document the range of localization patterns observed across the analysed specimens, including both positive and signal-negative fields. Because DiD is a lipophilic membrane dye that can be progressively diluted during cell division and reduced through membrane turnover, later observations reflect persistence of detectable fluorescence rather than an absolute measurement of transplanted-cell survival, abundance, proliferation, or differentiation.

**Figure S10:**
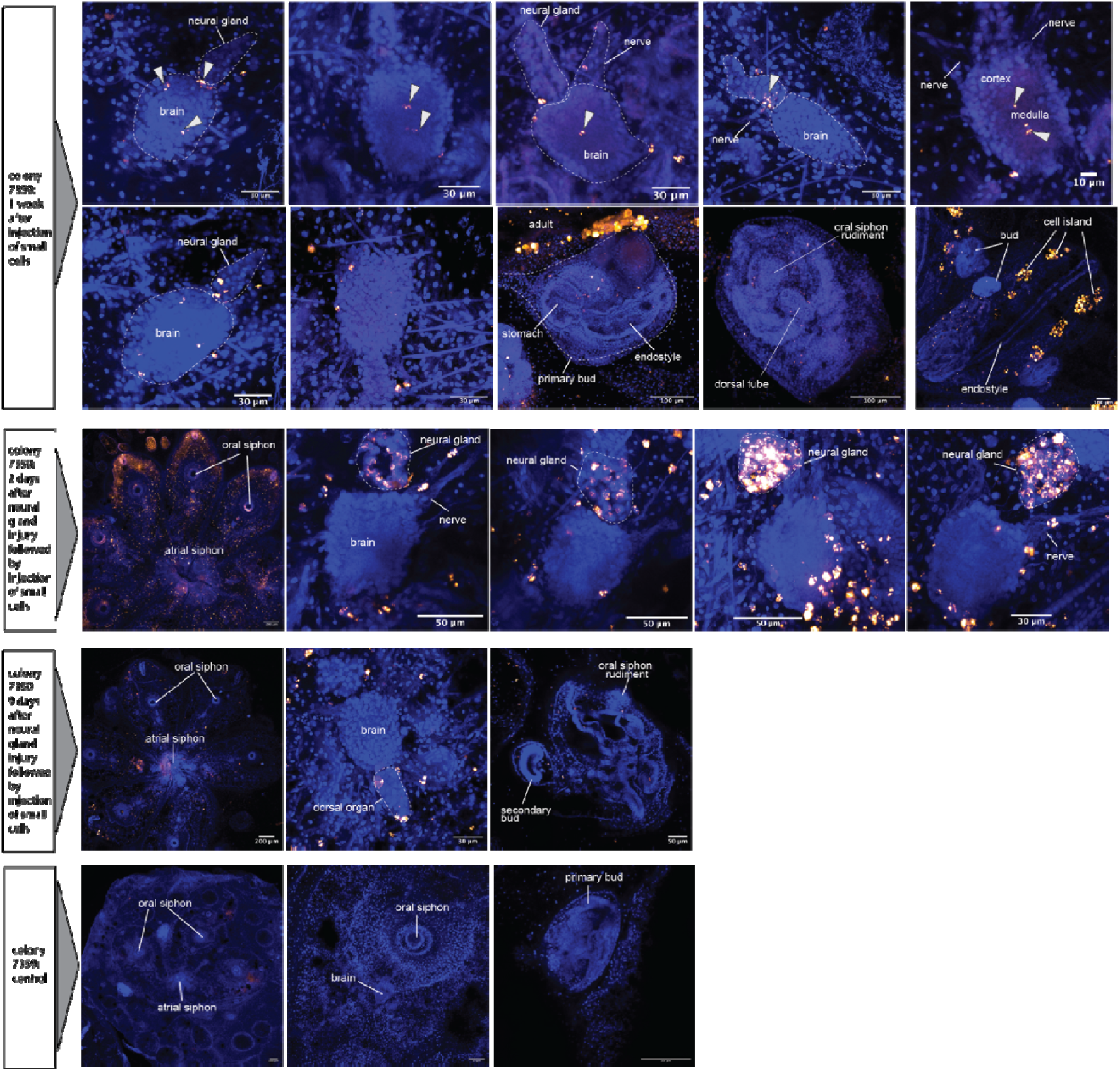
Confocal tracking of injected DiD-labelled small cells in adult and developing neural-complex-associated territories. Representative confocal images showing the distribution of DiD-labelled small cells after in vivo transplantation into recipient *Botryllus schlosseri* colonies. Nuclei are shown in blue and DiD-labelled cells or cell aggregates are shown in red/orange. In uninjured colonies examined approximately one week after injection, labelled cells were detected within and around the adult brain, at the brain–neural gland interface, along peripheral nerve-associated regions, and in developing bud-associated territories. Dashed outlines indicate the approximate boundaries of the brain and neural gland where shown, and arrowheads indicate representative DiD-positive cells or aggregates. Additional whole-zooid and bud-level views document labelled cells in adult tissues, the primary bud, the secondary bud, dorsal tube-associated regions, and cell-island/endostyle-associated territories. Following targeted injury to the neural gland region, numerous DiD-labelled cells were observed around the neural gland and adjacent brain one day after injury. Labelled cells remained detectable in neural-complex-associated regions nine days after injury, including around the neural gland, brain, and nearby nerves. Images are representative of the available confocal dataset and were included to document the range of anatomical localizations observed across specimens. Scale bars are indicated in each panel. Because DiD fluorescence can be progressively diluted during cell division and can decrease with membrane turnover over time, the images indicate persistence of detectable label rather than an absolute measure of transplanted-cell survival, proliferation, or integration.

